# Carbon bias from grassy tree misclassification: revealing forest structural heterogeneity by integrating GEDI and Sentinel-2

**DOI:** 10.64898/2025.12.25.696531

**Authors:** Aiyu Zheng, Yi Yin, Mingzhen Lu

## Abstract

Tropical forests store roughly half of terrestrial carbon, yet carbon estimates in regenerating and disturbed forests remain highly uncertain. A major source of bias is the prevalence of fast-growing, canopy-forming monocots—such as bamboo, palms, and bananas—that are often misclassified as trees. These “grassy trees” achieve canopy dominance but lack secondary growth, violating woody allometries used in most biomass models. Although NASA’s GEDI mission has transformed large-scale biomass mapping with spaceborne LiDAR, its products rely on coarse plant functional types (PFTs), causing grassy-tree-dominated canopies to be absorbed into evergreen broadleaf tree (EBT) classes. Using a texture-based Sentinel-2 classifier, we isolated bamboo-dominated forests within GEDI EBT products in Xishuangbanna, China. GEDI observations show that bamboo canopies are structurally distinct from tree-dominated forests and lead to systematic carbon overestimation of 20–44 Mg C ha⁻¹ relative to empirical benchmarks. Our framework improves carbon accounting in structurally heterogeneous forests while remaining adaptable for place-based management.

## 1. Introduction

Tropical forests are a crucial component of the global carbon cycle, storing approximately 50% of terrestrial carbon. While old-growth forests hold large carbon stocks and biodiversity, regenerating and frequently disturbed forests have expanded from 16.2% to 28.9% of total tropical forest area over the past 30 years, sequestrating carbon at rates five times higher than intact forests (Pan et al. 2024). However, their carbon dynamics remain highly uncertain (±20%) despite their increasing importance in land carbon sinks. This uncertainty is amplified in regrowing forests because disturbance and recovery generate fine-scale structural and compositional heterogeneity that challenges mapping and biomass inference.

A prominent contributor to this heterogeneity is the prevalence of canopy-forming monocots—large monocots such as bamboo, palms, and bananas—whose canopy architecture and biomass allocation systematically depart from the tree-dominated canopies that plant functional type (PFT) labels like “evergreen broadleaf trees” implicitly represent. These “grassy trees” have reached tree-like canopy dominance but are constrained by the lack of secondary growth like grasses with densely packed herbaceous tissues, often have hollow stems and lower aboveground biomass density than typical trees (Avalos et al. 2022; Zheng and Lu 2025). They can dominate disturbed tropical canopies (e.g., bamboo covering up to 21.3% of southwestern Amazonian forests; (de Carvalho et al. 2013) and may sequester carbon at rates several times higher than typical regrowth forests (Heinrich et al. 2021; Nath et al. 2015). Differences in above- and belowground biomass of grassy trees (Yuen et al. 2017; Navarro et al. 2008; Kotowska 2015; Song et al. 2017) further complicate tree-based biomass estimations where they are abundant.

Recent advancements in high-resolution remote sensing offer a path forward in addressing these challenges arising from complex canopy structures associated with grassy tree abundance in disturbed and fragmented landscapes. Sentinel-2 from the European Space Agency’s Copernicus programme (10-m; global since 2015; (Drusch et al. 2012)), coupled with texture-based analysis techniques, provides new opportunities to identify forest areas dominated by bamboo and palms. Unlike traditional spectral classification, texture-based approaches such as the Gray-Level Co-occurrence Matrix (GLCM) capture canopy structural patterns that distinguish grassy tree canopies from those formed by broadleaf woody trees (Humeau-Heurtier 2019). For example, sub-meter-resolution Maxar imagery shows that grassy trees such as bamboo form elongated or patchy canopy patterns, in contrast to the rounded crowns of broadleaf woody trees (see Fig.1 and Fig.A.1).

**Fig 1.**
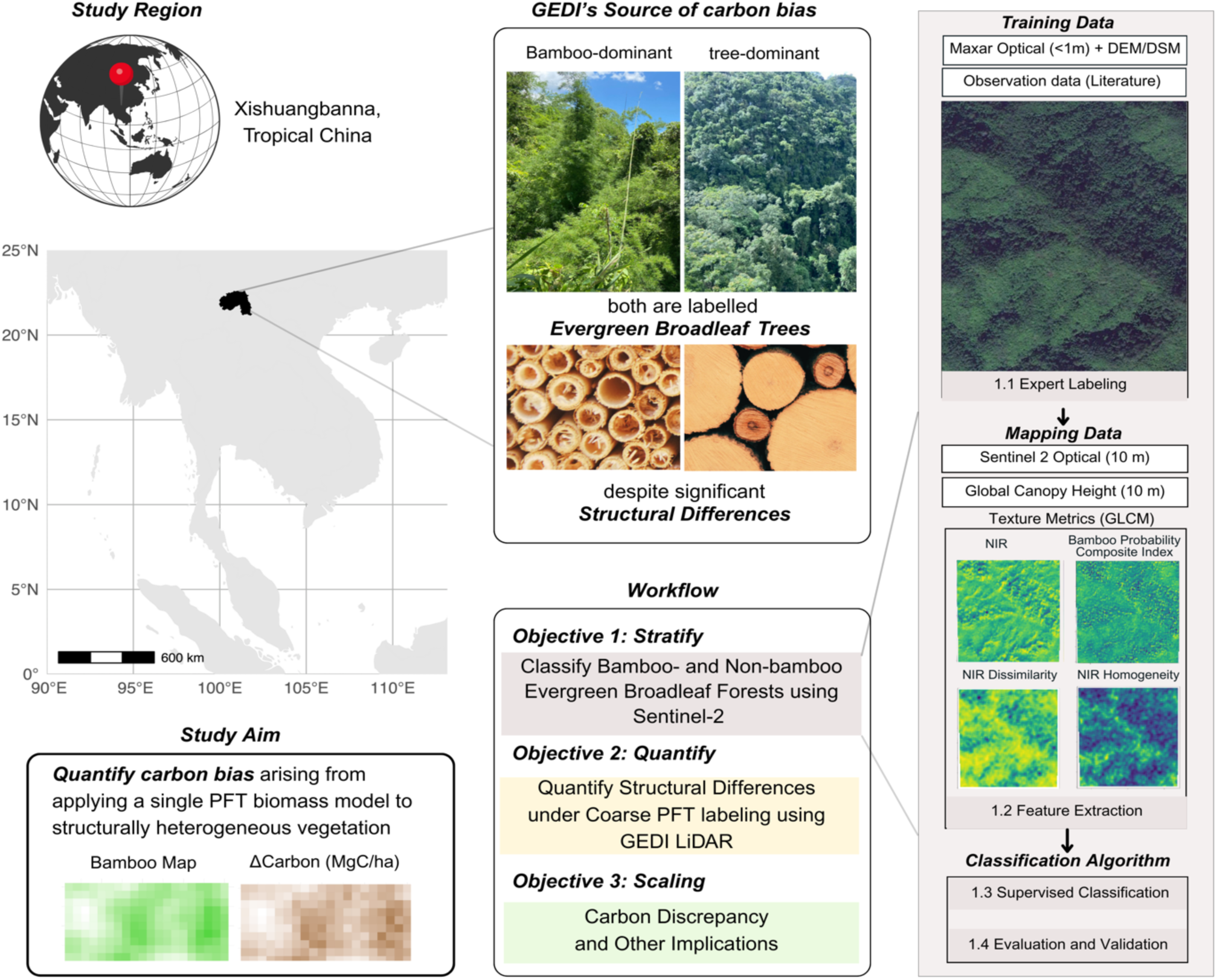
Conceptual workflow of this study, combining texture cues from Sentinel-2 with vertical structure from a global canopy height model (Lang et al. 2022) and GEDI LiDAR metrics to quantify carbon bias in functionally heterogeneous evergreen broadleaf tree (EBT) forests with high bamboo abundance in Xishuangbanna.

These canopy texture cues provide the horizontal context that complements vertical insights from LiDAR missions such as NASA’s Global Ecosystem Dynamics Investigation (GEDI), which delivers global products for canopy height (L2A), canopy cover and vertical structure (L2B), and aboveground biomass density (L4A) (Dubayah et al. 2022). These structural insights are crucial to address two primary barriers that have prevented the integration of grassy trees into forest biomass assessment in previous research. First, forest inventories have historically overlooked these groups due to ambiguous growth forms, leading to underrepresentation in ground-truth data (Fadrique et al. 2020). Second, remote sensing has struggled to accurately identify these functionally unique species within forest canopies. Grassy trees share spectral properties similar to those of broadleaf trees (Li et al. 2019; Yusof et al. 2021), while lower-resolution sensors such as Landsat (30 m) and MODIS (250 m) lack the spatial detail needed to detect them (Qi et al. 2022; Venkatappa et al. 2020; Jia et al. 2020). This spectral and spatial ambiguity has compounded existing ground-based biases, reinforcing uncertainties in forest carbon estimates.

However, GEDI LiDAR products have limitations because they stratify global forests into five PFTs—evergreen broadleaf trees (EBT), deciduous broadleaf trees, evergreen needleleaf trees, deciduous needleleaf trees, and a combined grass–shrub–woodland class—informed by MODIS land cover inputs (Kellner et al. 2023). While suitable for global-scale modeling, this coarse classification can obscure critical structural differences among vegetation subtypes, especially in regions with high functional heterogeneity. Applying a single PFT label (e.g., EBT) across diverse canopy structures—such as tree-dominated, palm-rich, or bamboo-dominated forests—can introduce systematic biases in GEDI-based biomass estimates (Bruening et al. 2023; Lin et al. 2025). These issues are particularly acute in tropical Asia and Africa, where calibration data are sparse, and canopy forms often depart from the tree-based assumptions embedded in GEDI biomass models (Dubayah et al. 2022).

To address these limitations, we combine GEDI with complementary datasets to improve grassy tree stratification and quantify carbon-estimation bias from mis-labeling grassy trees as EBT. While GEDI provides high-fidelity vertical structure metrics, it lacks the horizontal context needed to disaggregate compositionally diverse forest types. Sentinel-2, despite its limited sensitivity to vertical structure, adds valuable context through its consistent spatial resolution and spectral–textural richness. When paired with texture-based approaches, it can reveal patterns that help distinguish vegetation types otherwise lumped under the same PFT (Humeau-Heurtier 2019; Descals 2023).

We conceptualize this integration in Fig.1, where Sentinel-derived compositional context helps subdivide GEDI’s broad PFT class, EBT, into grassy trees (bamboo) and woody trees, identifying areas where structural and biomass deviations are likely and thus require calibration refinement. We then build a specific sub-PFT (e.g., bamboo) classifier for mapping that is designed to generalize. Together, Fig.1 presents the core workflow of this study—combining structure and composition via texture-enhanced satellite imagery and GEDI LiDAR—to stratify heterogeneous forests more accurately and quantify functional misclassification biases in carbon accounting.

In this study, we focus on bamboo-dominated forests in Xishuangbanna, southwestern China—a region with high bamboo richness, natural abundance, and cultivation, yet currently subsumed under GEDI’s evergreen broadleaf tree (EBT) class. We use this region to demonstrate how combining texture-enhanced Sentinel-2 classification with GEDI structural metrics refines carbon assessments and maps structurally distinct sub-PFT canopies. While this study centers on bamboo, the workflow is designed to be extended to other grassy-tree-dominated systems such as palm forests, as well as other structurally heterogeneous forests, like those with high liana abundance. Specifically, our objectives are outlined in Fig.1:

1. **Stratify** bamboo-dominated forests from tree-dominated EBT forests and other land-use types using a texture-enhanced Sentinel-2 classification.
2. **Quantify** structural and biomass differences in GEDI metrics between classified bamboo and non-bamboo EBT forests to evaluate bias from GEDI’s tree-centric EBT biomass model.
3. **Scale** aboveground biomass density to aboveground carbon density and map the spatial pattern of carbon bias in forests containing sub-PFT bamboo canopies.

## 2. Methods

### 2.1 Study Region

Xishuangbanna, on the northern edge of tropical Asia, hosts true tropical rainforests despite its monsoonal climate (Zhu et al. 2006). A biodiversity hotspot with over 5,000 higher plant species in just 19,690 km², the region has experienced extensive land-use change over recent decades (Li et al. 2008). Forest cover has declined from ∼60% in the 1950s due to the expansion of rubber plantations, cash crops, and shifting cultivation (Zhang and Cao 1995; Senf et al. 2013)(Zhang and Cao 1995); (Zhang and Cao 1995; Senf et al. 2013), giving rise to widespread secondary vegetation—especially bamboo forests (*Dendrocalamus membranaceus*, *Cephalostachyum pergracile*, etc., Yang et al. 2008)—now common along forest edges and within fragmented landscapes (bamboo in Fig.1).

Xishuangbanna presents an ideal setting to develop and evaluate our method, given its highly heterogeneous forest matrix, strong bamboo presence, and the cultural significance of bamboo among the local Dai population (Wang et al. 2002). We focus on this region to assess how the broad classification of bamboo as evergreen broadleaf trees in GEDI products can affect biomass estimation and carbon accounting.

### 2.2 Remote Sensing and Analytical Workflow

To accomplish the three objectives of our study, we designed an integrated remote sensing and analytical workflow that includes training data compilation, texture analysis, classification and accuracy assessment, evaluation of GEDI’s limitations, and the quantification of carbon discrepancies (Fig.1). Below, we describe each component of this workflow in detail.

#### 2.2.1 Datasets and Preprocessing

We aligned GEDI footprints to the 10-m Sentinel-2 grid by assigning each footprint to the 10-m pixel containing its geolocation (footprint center). Land-cover labels were extracted at that pixel and linked to footprint-level GEDI metrics for subsequent analyses.

##### Sentinel-2 (Optical, 10 m resolution)

We obtained Sentinel-2 imagery from Google Earth Engine (COPERNICUS/S2_harmonized) and generated four seasonal composites (January–March, April–June, July–September, and October–December) for the year 2021, selecting scenes with less than 5% cloud cover. Each composite included nine spectral bands (B2–B8, B11, B12) and was processed using QA60 and s2cloudless cloud masks to reduce atmospheric noise.

##### The Global Canopy Height Model

(10 m, Lang et al. 2022): a probabilistic deep learning model that fuses sparse height data from GEDI LiDAR with dense optical satellite images from Sentinel-2.

##### GEDI (LiDAR, 25 m resolution)

GEDI datasets provided canopy height and relative height metrics (Level 2A), canopy cover and vertical structure (Level 2B), and aboveground biomass density (Level 4A). All GEDI footprints (25-m resolution) were aligned to 10-meter Sentinel-2 composites for value extraction and analysis.

#### 2.2.2 Training data

We compiled a training dataset encompassing diverse vegetation and land-use categories, including bamboo, tree-dominated evergreen forests, rubber plantations, fruit and tea plantations, farms, construction areas, and water bodies. Labels were manually interpreted from 2021 Google Earth imagery (licensed Maxar submeter imagery), chosen for its minimal cloud cover and alignment with GEDI and Sentinel-2 temporal windows.

We reliably identified bamboo based on its distinct texture and canopy structure (Fig.1), supported by expert knowledge of its landscape distribution—such as along riparian corridors, within rubber plantations, and near forest edges—and by regional vegetation accounts (Zhu 2006; Zhu et al. 2015) describing *Dendrocalamus membranaceus* forests. We identified other vegetation types using a combination of phenological signals (e.g., seasonal reflectance in rice paddies), spatial context (e.g., farms near villages), auxiliary map labels (e.g., “Xishuangbanna Tropical Botanical Garden”), and literature-based spatial descriptions (Min et al. 2019; Li et al. 2008), drawing from multiple lines of evidence.

In Appendix A, we summarized the training data’s sample size and geographic coverage in Table A.1 and Fig A.1 and detailed the visual interpretation criteria used for each land-use type in Table A.2. Fig A.2 shows the strong spectral separability of the October-September composite for Xishuangbanna in 2021.

#### 2.2.3 Feature Extraction

For each labeled polygon, we extracted a comprehensive suite of features that capture spectral, textural, and contextual information (Fig. A.3). *Spectral features* include surface reflectance from selected Sentinel-2 bands (e.g., red, red-edge, SWIR), along with vegetation indices that characterize greenness, moisture, and phenology: Normalized Difference Vegetation Index (NDVI = (NIR – Red)/(NIR+ Red)), and the Bamboo Phenological Characteristic Index (BPCI = Red Edge 1/NIR-NIR/SWIR 1) (Huang et al. 2024). These features were computed from the seasonal composite imagery described earlier.

#### 2.2.4 Texture Metrics

We applied Gray-Level Co-occurrence Matrix (GLCM) techniques to characterize the spatial distribution of pixel intensities, capturing distinct texture patterns associated with bamboo’s unique leaf morphology and clumping architecture. Metrics such as pixel dissimilarity, homogeneity, and entropy have proven useful in distinguishing different species within mixed canopies (Mohammadpour et al. 2022). To identify the most informative spectral regions, we examined how texture metrics varied across different vegetation types—especially between bamboo-, tree-dominated forests, and rubber plantations. This analysis reduces dimensional redundancy and minimizes overfitting risks by selecting texture features (Band 8 dissimilarity and B8 homogeneity computed using a 5×5-pixel sliding window) that are less correlated yet discriminative.

#### 2.2.5 Classification algorithm and accuracy assessment

We trained and evaluated two machine learning classifiers: Random Forest (RF) and Convolutional Neural Networks (CNN) (detailed in Appendix B). RF is robust to noise and well-suited for integrating heterogeneous feature sets, while CNN enables pixel-based classification with spatial context sensitivity. To compare their performance, we withheld a subset of ground-truth data and applied both classifiers independently.

To assess classification accuracy, we conducted visual cross-validation by randomly sampling 100 points within each land-use class and verifying their true identity in Google Earth Pro (v7.3.6, Google Earth 2025) using high-resolution imagery from 2020 to 2022. Confusion matrices summarizing the results of this photo-interpretation-based validation are presented in Table C.1. We report standard accuracy metrics for the better-performing RF classifier: overall accuracy (the proportion of correctly classified cases), class-wise sensitivity (recall; true positive rate) and specificity (true negative rate), and the F1 score (which penalizes both missed detections and false positives).

#### 2.2.6 GEDI overlay

To evaluate the limitations of GEDI-derived products in distinguishing bamboo-dominated forests, we overlaid GEDI Level 2A (canopy height), Level 2B (canopy cover and vertical structure), and Level 4A (aboveground biomass) with our classified land cover map (Figure C.1). For each GEDI footprint intersecting a bamboo-classified pixel, we extracted its associated canopy height, structural, and biomass metrics. For comparison, the same set of variables was extracted from footprints located within tree-dominated forests and other land-use classes.

To account for classification uncertainty—particularly confusion between bamboo and tree categories—we treated the validation-derived misclassification rates as an observation model (confusion matrix; Table C.1) and applied a Bayesian measurement-error correction to infer the posterior true area of each land-cover class (detailed in Appendix C). This yields classification-error-adjusted estimates of the total extent of bamboo-dominated forests and other vegetation types across Xishuangbanna.

At the footprint level, we propagated this classification uncertainty by assigning each GEDI footprint a posterior probability of belonging to bamboo versus tree conditional on its mapped label, and then reweighted footprint-level summaries accordingly. This probabilistic reweighting enables more robust estimation of biomass distributions for bamboo- and tree-dominated forests and quantifies the extent to which biomass estimates are biased when bamboo-dominated canopies are treated as evergreen broadleaf forests under the default GEDI L4A EBT biomass model.

#### 2.2.7 Aboveground biomass bias from GEDI EBT model

GEDI Level 4A aboveground biomass density (AGBD, Mg ha⁻¹) estimates in Xishuangbanna are derived from a global EBT biomass model parameterized on GEDI relative height (RH) metrics (Kellner et al. 2023):

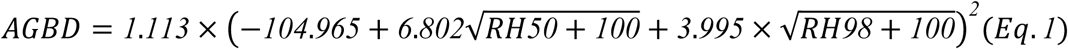

RH metrics summarize the vertical structure of a LiDAR footprint as percentiles of the cumulative GEDI waveform energy returned from height *z*. RH98 approximates the upper canopy envelope (near-maximum vegetation height), whereas RH50 represents the median height of returned vegetation material, capturing the vertical distribution of canopy structure.

However, this EBT model is applied uniformly across evergreen broadleaf forests, including bamboo-dominated stands that differ fundamentally from trees in growth form and vertical structure. To test whether this unstratified application induces systematic overestimation and to quantify its effect size, we (1) compared GEDI Level 2A/2B structural metrics between bamboo- and tree-dominated areas classified by our random-forest land-cover map, and (2) used empirical bamboo biomass benchmarks from the same ecoregion (Yuen et al. 2017) as external references for bias quantification (see Table D.1). Given strong management and land-use contrasts, we further separated bamboo into forest versus fallow contexts to obtain context-specific benchmarks and bias estimates.

### 2.3 GEDI structural comparison

We extracted GEDI Level 2A and 2B metrics from footprints whose surrounding area was classified as bamboo or non-bamboo EBT forests (trees). We compared their canopy height (RH98), relative height profile, plant area index (PAI), and plant area volume index (PAVD) throughout the canopy, to reveal structural differences that are masked when all these forests are labelled as a single EBT class in GEDI-based products. These structural contrasts motivate subsequent bias analysis.

#### Empirical benchmarks for bamboo aboveground biomass and carbon density

Established methods for estimating individual bamboo biomass often are typically based on diameter and height of clumpy tropical bamboos (Xayalath et al. 2019; Camargo García et al. 2023; Huy et al. 2019). Unlike trees, bamboo culms do not exhibit continuous secondary growth, and stand-level biomass depends strongly on age structure and culm turnover rather than height alone (Yen 2016). These demographic properties cannot be retrieved from GEDI waveform metrics. Our objective is therefore not to recalibrate bamboo allometry, but to evaluate whether a tree-based RH model produces systematic overestimation on bamboo and to quantify its effect size relative to empirically observed ranges.

Accordingly, we used empirical aboveground carbon density estimates synthesized by (Yuen et al. 2017) for tropical bamboos in Indochina (Myanmar, Laos, Thailand, Vietnam, and India), a region biogeographically adjacent to Xishuangbanna and characterized by comparable species composition. Because Xishuangbanna and Indochina share a contiguous Indo-Burma floristic context with overlapping bamboo taxa and broadly similar monsoonal tropical–subtropical climates (Zhu et al. 2006), we expect these synthesized carbon-density benchmarks to be broadly transferable to bamboo-dominated stands in our study region.

Reported summary statistics (mean, standard deviation, and max) were used to derive confidence intervals for comparison with GEDI L4A footprint-level estimates (detailed in Appendix D). To reflect agroforestry mosaics abundant with secondary forests in Xishuangbanna, bamboo pixels were classified as forest or fallow based on a 3×3 neighborhood rule: bamboo cells surrounded by ≥ 5 bamboo or tree neighbors were classified as forest bamboo; otherwise as fallow bamboo.

For bamboo footprints, we compared GEDI L4A AGBD estimates with empirical bamboo benchmarks stratified by bamboo context *i* ∈ {*forest, fallow*}, and quantified aboveground carbon bias as:

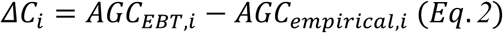

where aboveground carbon density (AGC) was derived by applying a constant carbon fraction (CF = 0.5) to AGBD for both bamboo and trees. Although CF varies among species and tissues, it has been inconsistently treated across the bamboo literature, whereas CF = 0.5 is widely used as a default in forest carbon accounting (Pan et al. 2011). We therefore adopt a single CF to ensure comparability between GEDI-derived AGBD and empirical benchmarks, noting that alternative defaults (e.g., CF = 0.47; IPCC 2006) would linearly rescale *ΔC* without affecting its sign. Positive *ΔC* values indicate overestimation of bamboo carbon by GEDI’s EBT model relative to empirical benchmarks.

##### Spatial extrapolation of carbon bias across Xishuangbanna

To characterize the spatial distribution of aboveground carbon bias (*ΔC*) across bamboo-rich landscapes in Xishuangbanna, footprint-level estimates must be extrapolated to continuous pixel-based surfaces. To ensure internal consistency with GEDI’s operational biomass products, this extrapolation was conducted entirely within the framework of the GEDI EBT biomass model (Eq. 1). Because the EBT model requires both RH50 and RH98 as inputs, we first quantified the statistical relationship between RH50 and RH98 separately for bamboo and tree footprints using GEDI Level 2A data. We applied the class-specific RH50–RH98 relationships to the 10-m global canopy height model (Lang et al. 2022) as an operational approximation to enable internally consistent regional mapping within our framework. Because the Lang et al. (2022) product is generated by fusing sparse GEDI Level 2 RH98 observations with dense Sentinel-2 optical data, it provides a GEDI-anchored proxy for canopy stature; nevertheless, it is not a one-to-one substitute for footprint-level RH98. Any residual mismatch between this 10-m canopy height layer and GEDI RH98 over overlapping footprints in Xishuangbanna is therefore treated as a limitation of our extrapolation. The estimated RH50 and RH98 were then used to compute pixel-level AGBD and aboveground carbon density using the EBT biomass model (Eq. 1).

The analytical form of the EBT biomass model (Eq. 1) does not capture all calibration steps and ancillary corrections used to generate official GEDI Level 4A biomass estimates. To quantify the systematic error introduced by applying this simplified RH-based formulation alone, we compared AGBD estimates derived from Eq. 1—using RH metrics from GEDI Level 2 data—with published Level 4A AGBD values for footprints where Level 2 and Level 4 observations overlapped.

Finally, we applied Eq. 1 across the entire Xishuangbanna region and converted AGBD to aboveground carbon density. To illustrate the spatial magnitude of overestimation, carbon bias was expressed as a percentage of EBT-estimated carbon density and mapped separately for forest and fallow bamboo areas:

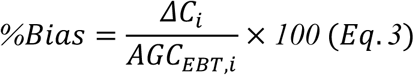

This multi-step extrapolation introduces three main sources of uncertainty: (1) misclassification errors from the random-forest land-cover map, (2) uncertainty in the class-specific RH50–RH98 regression models, and (3) systematic error arising from applying the simplified EBT biomass formulation (Eq. 1). These uncertainties were explicitly evaluated and propagated in subsequent analyses (see Results).

## 3. Results

### 3.1 Reliable sub-PFT land-cover classification enables robust bamboo detection

The random forest classifier achieved strong overall performance in distinguishing bamboo from other land-cover classes across Xishuangbanna (Fig. 2). Feature distributions derived from spectral texture (Band 8—Near Infrared’s homogeneity and dissimilarity), canopy height, and the Bamboo Phenological Characteristic Index (BPCI) show clear separation among vegetation classes in the training data (Fig. 2a), indicating that these features capture complementary structural and phenological information relevant for bamboo detection.

**Fig 2.**
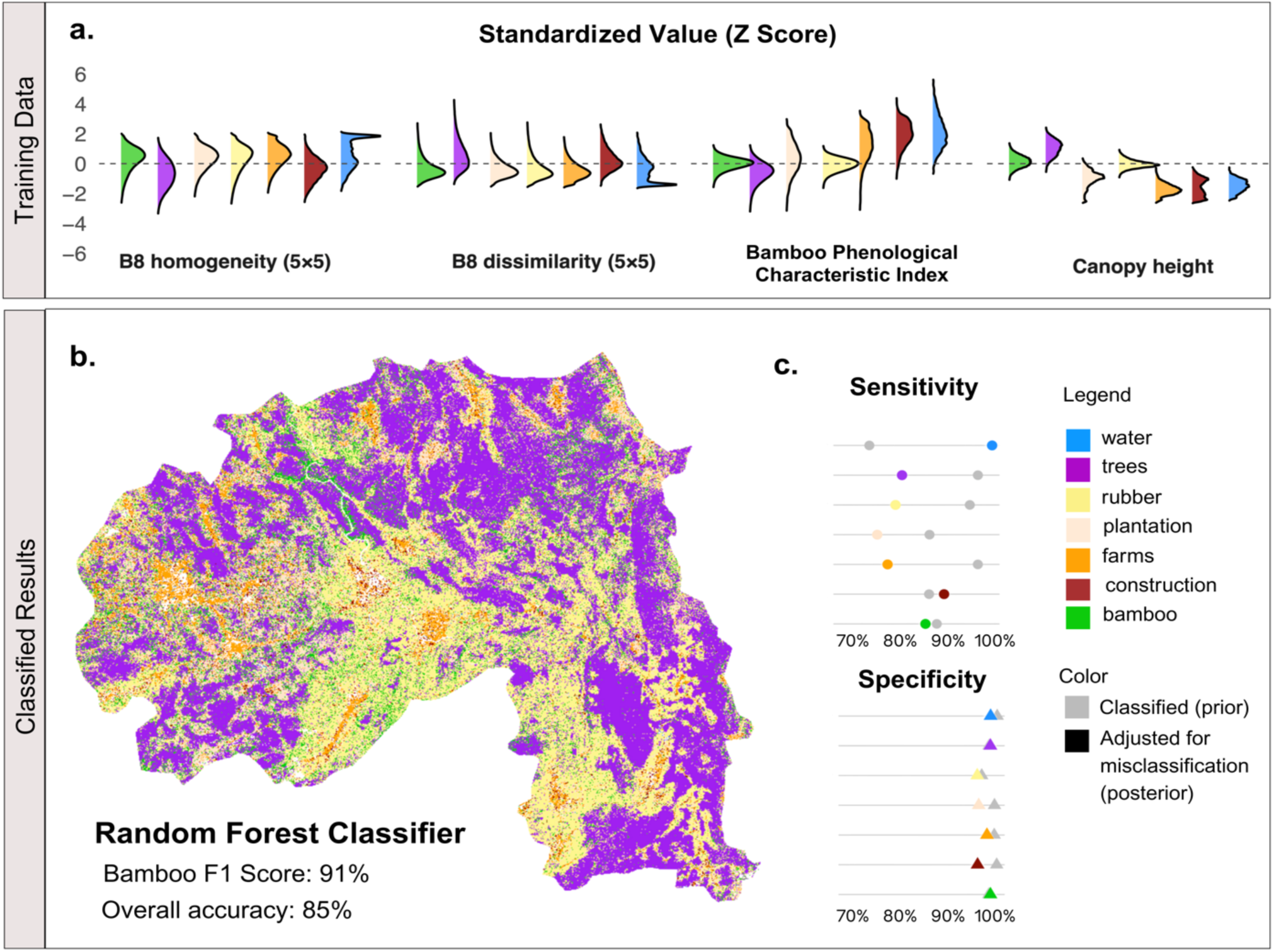
Sub-PFT mapping reveals hidden vegetation heterogeneity in Xishuangbanna. **a**) Standardized distributions (Z-scores) of four key predictors in the training data—Band 8 (near-infrared) homogeneity and dissimilarity (5×5-pixel window), the Bamboo Phenological Characteristic Index, and canopy height—show that the seven mapped classes occupy distinct regions of feature space, indicating that the training data and selected predictors provide strong discriminatory power for subsequent classification. **b**) Using Sentinel-2 spectral indices and GLCM texture features, we classified sub-PFT vegetation and land-cover types across Xishuangbanna with a Random Forest model. The resulting 10-m map reveals fine-scale mosaics of bamboo patches embedded within forest, rubber, and agricultural matrices—heterogeneity typically masked by the EBT forest label in GEDI-based products. **c**) Class-wise sensitivity (circles) and specificity (triangles) estimated from 70/30 hold-out validation during training (grey symbols) and from visually interpreted sample points drawn from the classified map (colored symbols), which we used to adjust for misclassification. Bamboo achieved an F1 score of 0.91 (overall accuracy 0.85), enabling reliable downstream quantification of bamboo cover and configuration.

The resulting land-cover map reveals spatially coherent patterns of bamboo, trees, rubber plantations, and agricultural land consistent with known landscape organization in the region (Fig. 2b). For instance, bamboo-dominated riparian belts are apparent along the Lancang River corridor, consistent with reported patterns of river-associated bamboo stands in the region. Beyond riparian zones, bamboo also occurs as patchy elements embedded within rubber plantations and farmland mosaics, consistent with local smallholder management practices. These spatial patterns are further supported by high-resolution aerial maps (Fig A.1).

Classification performance was strong despite the highly fragmented tropical landscape, with an overall accuracy of 85% and a bamboo F1 score of 91% (Fig. 2b), indicating reliable bamboo detection with low omission. Nonetheless, confusion among spectrally and structurally similar vegetation classes—most notably bamboo, trees, and rubber plantations—remained non-negligible, consistent with frequent co-occurrence and fine-scale mixing of these vegetation types in agroforestry mosaics. Class-specific sensitivity and specificity (Fig. 2c) further indicate that residual misclassification persists even under a well-performing classifier. We therefore explicitly propagated this classification uncertainty in subsequent analyses using a Bayesian posterior resampling framework and Monte Carlo reassignment of class labels, rather than treating the classified map as error-free. The posterior updating methods and results are detailed in Appendix C.

### 3.2 GEDI reveals systematic structural differences between bamboo- and tree-dominated forests

Building on the validated land-cover classification and explicitly quantified uncertainty, we overlaid the classification map with GEDI LiDAR footprints to test whether bamboo- and tree-dominated forests exhibit systematic differences in canopy structure and biomass-related metrics as directly observed by GEDI.

To aid interpretation of subsequent results, Fig. 3a illustrates how GEDI LiDAR footprint-level observations characterize canopy structure. GEDI footprints sample forest structure within circular columns approximately 25 m in diameter, capturing the vertical distribution of vegetation from the ground to the top of the canopy. Within each footprint, the returned waveform resolves a vertical profile of vegetation material, allowing both canopy height and the allocation of plant material along the vertical axis to be directly observed.

**Fig 3.**
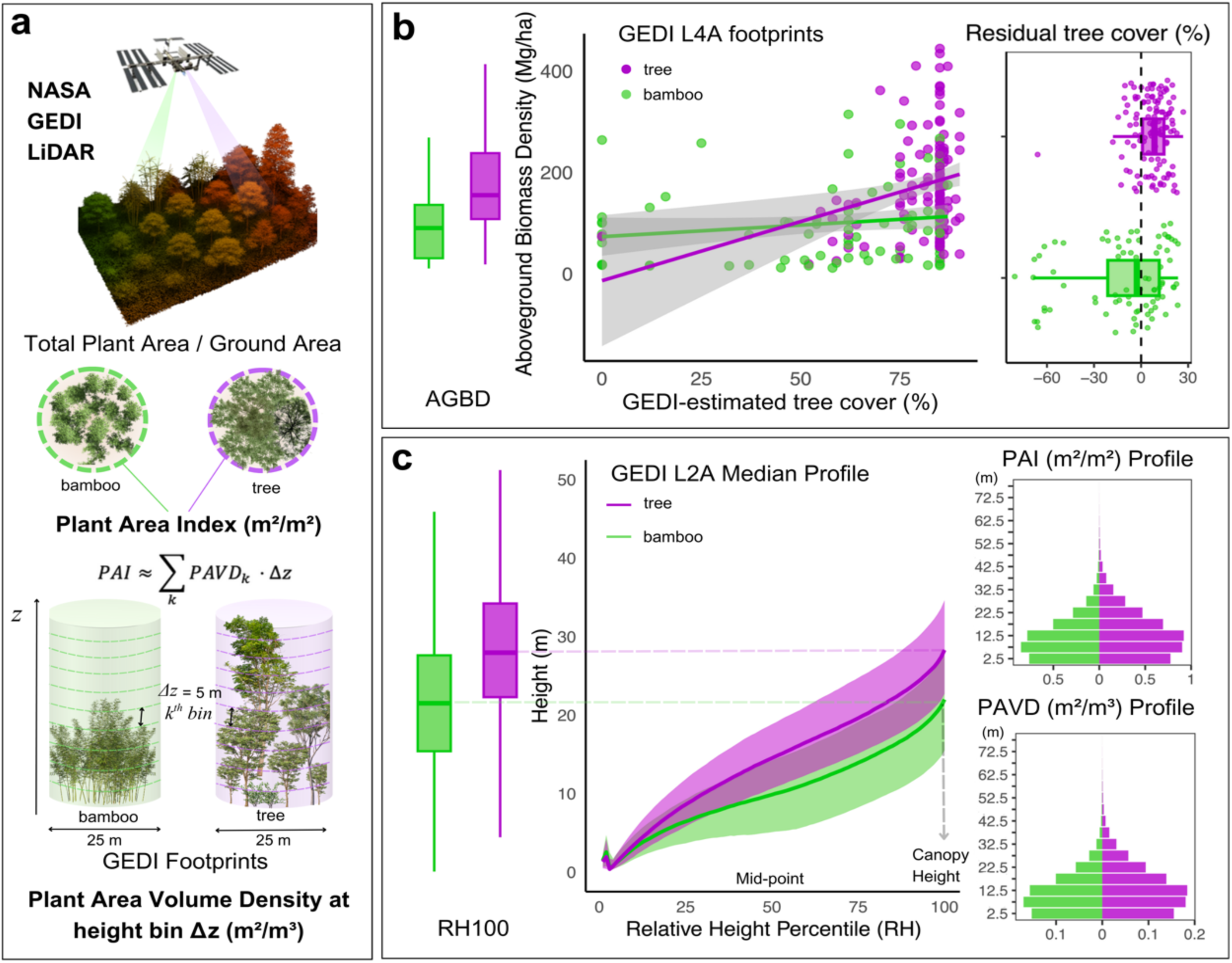
Structural differences between bamboo- and tree-dominated forests from GEDI. **(a)** Conceptual illustration of how GEDI waveforms capture vertical canopy structure within each ∼25 m footprint, from which canopy height, plant area index (PAI), and plant area volume density (PAVD) profiles are derived (see Wang et al. 2025). **(b)** Bamboo-dominated footprints have lower AGBD than tree-dominated footprints. Residual tree cover is defined as GEDI-reported tree cover minus the tree-cover value expected for a tree-dominated footprint with the same AGBD; values are consistently negative for bamboo, indicating its cover–biomass decoupling relative to tree-dominated forests. **(c)** Vertical structure differs strongly by vegetation type: bamboo has lower canopy heights (RH98) and a lower upper-canopy profile than trees, with broadly overlapping lower–mid canopy layers. PAI and PAVD profiles are consistent with these canopy-structure contrasts.

As shown in Fig. 3a, the vertical distribution of plant material is summarized by plant area volume density (PAVD) across successive height bins (*Δz*), while plant area index (PAI) reflects the cumulative plant area per unit ground area within the footprint. Conceptually, PAVD describes how plant material is distributed with height, whereas PAI indicates how plant biomass accumulates vertically, analogous to leaf area index but incorporating both foliage and woody components detected by the LiDAR waveform. Together, these GEDI-derived quantities provide an intuitive framework for interpreting the structural contrasts between bamboo- and tree-dominated forests shown in Fig. 3b–c and Table 1.

**Table 1.** Summary of structural and biomass differences between bamboo- and tree-dominated evergreen broadleaf forests as observed by GEDI LiDAR. Raw footprint-level medians (Panel A) are contrasted with misclassification-aware median differences obtained from posterior Monte Carlo resampling (Panel B; 1,000 iterations). All differences are expressed as Trees − Bamboo, and the fraction of Monte Carlo iterations with *p* < 0.05 quantifies the robustness of each contrast to classification uncertainty.

| **Panel A. Raw GEDI observations (classified bamboo vs trees)** |  |  |  |  |  |
| --- | --- | --- | --- | --- | --- |
| Metric | n<br>(Bamboo) | Median<br>(Bamboo) | n<br>(Trees) | Median<br>(Trees) | $\Delta$ Median<br>(Trees –<br>Bamboo) |
| Aboveground Biomass<br>Density (Mg ha<sup>-1</sup>) | 81 | 90.29 | 121 | 154.41 | 64.12 |
| Canopy Height (m) | 31,067 | 21.87 | 123,142 | 28.39 | 6.52 |
| Plant Area Index (m<sup>2</sup><br>m<sup>-2</sup>) | 43,538 | 3.28 | 171,085 | 4.08 | 0.81 |
| Total Vegetation<br>Volume (m<sup>3</sup> m<sup>-2</sup>) | 43,508 | 0.74 | 171,056 | 0.90 | 0.15 |
| **Panel B. Misclassification-aware contrasts (posterior Monte Carlo; n = 1000 iterations)** |  |  |  |  |  |
| Metric | $\Delta$ Median (MC,<br>2.5–97.5%) | MC Median | MC $p < 0.05$ | | |
| Aboveground Biomass<br>Density (Mg ha<sup>-1</sup>) | 44.67–66.46 | 54.37 | 1.00 |  |  |
| Canopy Height (m) | 5.19–5.32 | 5.26 | 1.00 |  |  |
| Plant Area Index (m<sup>2</sup><br>m<sup>-2</sup>) | 0.64–0.67 | 0.65 | 1.00 |  |  |
| Total Vegetation<br>Volume (m<sup>3</sup> m<sup>-2</sup>) | 0.12–0.13 | 0.12 | 1.00 |  |  |

Despite being processed using the same EBT biomass model, bamboo- and tree-dominated GEDI footprints exhibited markedly different Level 4A aboveground biomass density patterns (Fig. 3b). We use residual tree cover as a diagnostic of structural inconsistency: for each footprint, we calculate GEDI-reported tree cover minus the tree cover expected at the same AGBD from the unstratified AGBD–tree-cover relationship (i.e., treating all EBT as structurally equivalent). In tree-dominated footprints, AGBD increased strongly with GEDI-estimated tree cover, which was predominantly high (> 50%), consistent with the model’s underlying assumption that canopy cover and vertical extent scale with woody biomass. In contrast, bamboo-dominated footprints showed substantially lower AGBD (median 42% lower) and a weak cover-biomass coupling: similar AGBD values occurred across both low (< 25%) and high (> 50%) tree cover. Accordingly, residual tree cover was predominantly negative for bamboo footprints and positive for tree footprints, indicating a systematic decoupling between cover and biomass in bamboo that arises from GEDI observations alone, prior to any comparison with external biomass benchmarks. Altogether, GEDI tree cover is not an equivalent proxy for biomass in bamboo-dominated EBT forests as it is in tree-dominated EBT forests.

GEDI Level 2A metrics further indicate systematic differences in vertical canopy structure between bamboo- and tree-dominated forests (Fig. 3c; Table 1). Bamboo footprints exhibited lower canopy height, with median canopy height ∼23% lower than tree footprints. Relative height (RH) profiles show that bamboo canopies concentrate vegetation material at lower heights, resulting in a more compressed vertical distribution, whereas tree canopies exhibit greater vertical extension, and more plant material allocated to upper canopy layers. Consistent with these patterns, bamboo footprints showed lower total PAI and total vegetation volume, and distinct PAVD profiles across height bins. Together, these GEDI-observed structural contrasts provide a mechanistic explanation for the divergent biomass patterns in Fig. 3b and underscore that tree-based structural predictors embedded in the EBT model do not adequately represent bamboo-dominated forests.

To assess the sensitivity of these structural contrasts and biomass differences to land-cover misclassification, we propagated classification uncertainty using posterior resampling and Monte Carlo simulations. Across all four GEDI-derived metrics, median differences between tree- and bamboo-dominated footprints remained consistently positive and statistically significant in all Monte Carlo iterations (proportion of iterations with *p* < 0.05 = 1.0; Table 2). Accounting for misclassification reduced the magnitude of the estimated differences relative to naïve estimates—for example, the median tree–bamboo difference in aboveground biomass density decreased from 64.1 Mg ha⁻¹ to 54.4 Mg ha⁻¹—but did not alter their direction or overall effect size. Similar patterns were observed for canopy height, plant area index, and total vegetation volume (Fig C.2). These results indicate that the observed structural differences are robust to plausible classification errors and are not driven by misclassification alone.

**Table 2.** Key models and uncertainty components used to extrapolate bamboo carbon bias across Xishuangbanna. (A) Class-specific RH50–RH98 regressions fitted to GEDI Level 2 footprints (bamboo vs. tree) and used to predict RH50 from a 10-m canopy height layer (treated operationally as RH98) for regional application of the EBT biomass model. (B) Systematic discrepancy between biomass/carbon estimated from the simplified RH-only EBT formulation (Eq. 1) and the published GEDI Level 4A carbon at footprints where Level 2 and Level 4 observations overlap, summarized as residual carbon (*C_est,i_* – *C_L_*_4*A,i*_). (C) Misclassification-aware GEDI estimates of bamboo carbon bias relative to empirical benchmarks, obtained by posterior Monte Carlo resampling that propagates uncertainty in whether mapped-bamboo footprints are truly bamboo (conditional on the mapped label) and recomputes GEDI–empirical mean differences for forest- and fallow-bamboo contexts. The “Total” component combines the misclassification-aware GEDI–empirical bias with the EBT-formulation discrepancy in Panel B to produce the bias estimates used for regional mapping. Panel C reports bias estimates conditional on footprints mapped as bamboo (false positives are removed); omission errors (false negatives) are not propagated (see Discussion).

| Panel A. Class-specific RH50–RH98 regressions |  |  |  |  |  |  |
| --- | --- | --- | --- | --- | --- | --- |
| Class | Model | $\beta_0$ | $\beta_1$ | n | R<sup>2</sup> | RMSE (m) |
| Trees | RH50 = $\beta_0 + \beta_1 \cdot \text{RH98}$ | -2.316 | 0.628 | 123,142 | 0.675 | 3.74 |
| Bamboo | RH50 = $\beta_0 + \beta_1 \cdot \text{RH98}$ | -3.872 | 0.649 | 31,067 | 0.745 | 3.35 |
| Panel B. Discrepancy between RH-only EBT re-estimation and published GEDI L4A carbon at overlapping footprints |  |  |  |  |  |  |
| Residual (Mg C ha<sup>-1</sup>): $\Delta C_{EBT} = C_{est} - C_{L4A}$; $C_{est} = 0.5 \times AGBD_{EBT}(\text{RH50}, \text{RH98})$ | | | | | | |
| Class |  | n | Mean residual |  | SD |  |
| Bamboo |  | 81 | 5.31 |  | 51.52 |  |
| Trees |  | 121 | 3.97 |  | 44.49 |  |
| Panel C. Misclassification-aware bamboo carbon bias relative to empirical benchmarks |  |  |  |  |  |  |
| Naïve (no-correction) mean bias (Mg C/ha): Bamboo Fallow: 43.7; Bamboo Forest: 19.9 |  |  |  |  |  |  |
| Bamboo context | Component | Mean bias (Mg C/ha) |  | Median | 2.5% | 97.5% |
| Fallow bamboo | posterior-resampled bamboo bias (MC) | 44.1 |  | 43.7 | 29.6 | 59.0 |
| Forest bamboo | posterior-resampled bamboo bias (MC) | 19.8 |  | 19.8 | 16.4 | 23.3 |
| Fallow bamboo | Total (MC + EBT discrepancy) | 49.4 |  | 49.0 | 34.8 | 64.2 |
| Forest bamboo | Total (MC + EBT discrepancy) | 25.1 |  | 25.1 | 21.7 | 28.6 |

### 3.3 Systematic carbon overestimation in bamboo forests by GEDI EBT models

Building on the GEDI-observed structural differences between bamboo- and tree-dominated forests (Fig. 3), we next quantified the magnitude of biomass and carbon bias arising from applying the EBT model to bamboo-dominated systems using empirical benchmarks. Bamboo-dominated footprints were stratified into forest and fallow contexts based on local neighborhood composition, reflecting the heterogeneous agroforestry landscape of Xishuangbanna (Fig. 4a).

**Fig 4.**
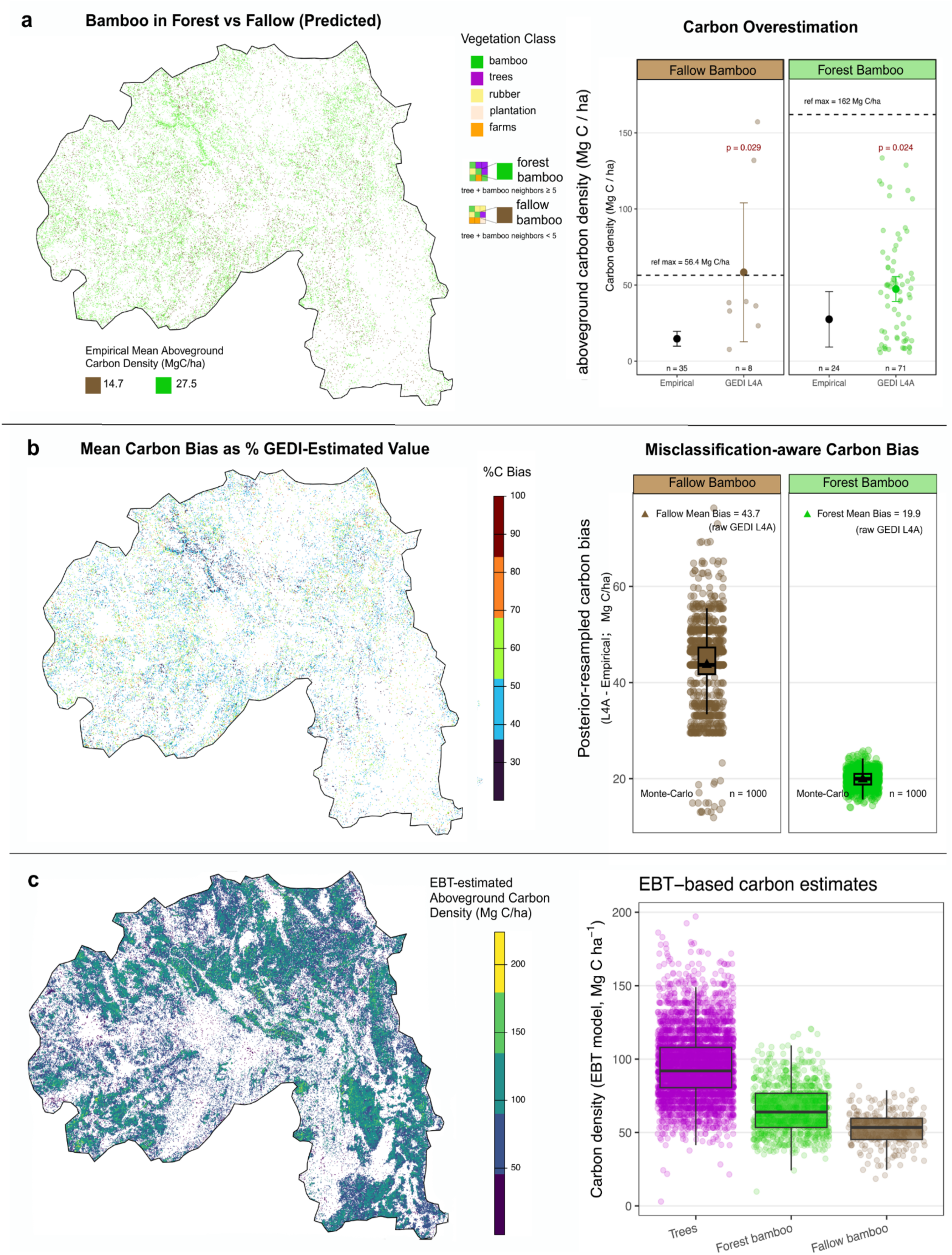
Quantifying carbon overestimation when structurally distinct evergreen broadleaf forests are treated with GEDI’s single PFT-based EBT biomass model in Xishuangbanna. **(a)** Predicted bamboo pixels were separated into forest versus fallow contexts using a 3×3 neighborhood rule (forest bamboo: ≥ 5 bamboo/tree neighbors; fallow bamboo: < 5). GEDI L4A aboveground carbon density for each bamboo type is compared with regional empirical reference values from Yuen et al. (2017). Error bars show t-based 95% confidence intervals for the group mean carbon density (mean ± t·SD/√n); p-values are from one-sided Welch’s t-tests (H₁: GEDI > empirical). A re-derivation of the literature summary statistics is provided in Appendix D. **(b)** Uncertainty arising from bamboo misclassification was propagated using Monte Carlo resampling, in which footprint-level “true” land-cover classes were repeatedly drawn from posterior probabilities derived from the random-forest classifier. These misclassification-aware bias distributions were combined with bias arising from GEDI’s simplified EBT biomass formulation based on RH98 and RH50 (Eq.1 applied in GEDI L4A for Xishuangbanna; Kellner et al., 2023) to estimate mean percent carbon overestimation across the region. Fallow bamboo exhibits larger absolute carbon bias. **(c)** Region-wide EBT-based aboveground carbon estimates derived from the simplified EBT RH model. The boxplots summarize the resulting EBT-based carbon densities for pixels classified as trees, forest bamboo, and fallow bamboo. Carbon uncertainties are summarized in Table 2.

Comparisons between GEDI Level 4A estimates and empirical bamboo benchmarks reveal systematic overestimation of aboveground carbon density in both bamboo contexts (Fig. 4a). GEDI-derived carbon densities for fallow bamboo were significantly higher than empirical values (one-sided Welch’s *t*-test, H1: GEDI > empirical, p = 0.029), and forest bamboo exhibited an even larger positive deviation from its corresponding benchmark (p = 0.024). To limit the disproportionate influence of extreme GEDI footprint values on summary statistics, we excluded outliers in GEDI L4A bamboo carbon density (forest vs fallow) using the Tukey rule (values outside 1.5 × IQR) prior to visualization and statistical comparison.

The magnitude of overestimation also differed between bamboo contexts: fallow bamboo showed higher absolute carbon bias (43.7 Mg ha⁻¹ versus 19.9 Mg ha⁻¹) than forest bamboo with greater carbon uncertainty, consistent with its lower canopy height and higher landscape complexity observed by GEDI (Fig. 3). In contrast, tree-dominated forests exhibited substantially higher carbon densities overall, as expected for woody evergreen broadleaf systems. Together, these results indicate that applying a single EBT model to bamboo-dominated forests inflates carbon estimates and obscures meaningful variation between bamboo forest and fallow systems.

Accounting for land-cover misclassification using posterior Monte Carlo resampling did not eliminate the inferred carbon bias in bamboo-dominated systems (Fig. 4b). Across 1,000 posterior resampling iterations, mean aboveground carbon bias remained consistently positive for both bamboo contexts. Fallow bamboo exhibited a mean overestimation of 44.1Mg C ha⁻¹, whereas forest bamboo showed a larger mean bias of 19.8 Mg C ha⁻¹, closely matching the naïve footprint-level estimates (triangles representing mean values). The narrow spread of the Monte Carlo distributions for forest bamboo indicates that plausible misclassification errors are insufficient to explain the observed magnitude of overestimation, while fallow bamboo with wider distributions suggests higher uncertainty in AGBD estimates likely due to mixed influences from rubber plantations and agroforestry mosaics. These results confirm that the positive carbon bias arises from structural mismatch between bamboo and the EBT biomass model rather than from classification uncertainty alone.

Having established that bamboo-specific carbon overestimation persists after explicitly accounting for classification uncertainty at the footprint level, we next examined how this bias propagates spatially across Xishuangbanna. To support regional extrapolation, we quantified class-specific relationships between RH50 and RH98 from GEDI Level 2 observations and evaluated the discrepancy between AGBD predicted by a simplified, RH-only EBT formulation and published GEDI Level 4A estimates. These components were combined to propagate uncertainty into regional estimates of absolute and percent carbon bias. Inputs to this propagation include (1) class-specific RH50–RH98 regressions, (2) the EBT–L4A biomass discrepancy, and (3) land-cover misclassification adjustments; the height dependence of the EBT structural discrepancy is diagnosed in Appendix E. Table 2 summarizes the regression equations and the full bias decomposition.

Mapping the percentage carbon bias revealed pronounced spatial heterogeneity across bamboo-dominated landscapes (Fig. 4b). Relative overestimation frequently exceeded 40% in bamboo-rich areas, with systematic differences between fallow and forest bamboo (Fig. 4c). Because percentage bias is defined relative to EBT-estimated carbon density (Eq. 3), spatial variation reflects the joint effects of context-specific absolute bias (*ΔC_i_*) and variation in the underlying EBT-estimated carbon baseline (*AGC_EBT,i_*). Overall, these spatial patterns reinforce the footprint-level inference that GEDI’s EBT model systematically overestimates bamboo aboveground carbon density, while the magnitude of overestimation varies predictably with bamboo stand context.

## 4. Discussion

### 4.1 Grassy trees drive carbon bias in tree-calibrated biomass models

Our results demonstrate systematic biomass and carbon bias in GEDI Level 4A products over evergreen broadleaf forests in Xishuangbanna, tropical China. This bias persists after Bayesian correction for bamboo–tree classification uncertainty, indicating that it arises primarily from structural mismatch between bamboo-dominated canopies and GEDI’s global EBT biomass model, rather than from misclassification alone. Understanding this bias therefore requires examining how the operational EBT model was trained and applied.

The GEDI EBT biomass model is trained on globally uneven data, with continental and Southeast Asia among the most underrepresented regions (Kellner et al. 2023). Although calibration relies on simulated GEDI waveforms linked to evergreen broadleaf forest field plots from 21 countries, only 326 simulated waveforms support the EBT stratum in South Asia, compared with 3,441 in South America and 834 in Africa (Kellner et al. 2023, Table 1). As a result, statistical support for EBT biomass estimation in tropical Asia is substantially weaker than in other regions, a limitation particularly relevant for Xishuangbanna at the northern margin of Southeast Asia.

Consistent with this limitation, the CEOS Land Product Validation framework recognizes that, where training data are sparse, biomass estimates must rely on models trained in other regions or plant functional types, implicitly assuming geographic and structural transferability (Duncanson et al. 2022; Kay 2021). While necessary for global coverage, this assumption introduces systematic bias when canopy structure and species composition deviate from those represented in training data, such as bamboo-dominated secondary forests relative to tree-dominated evergreen broadleaf forests.

Ecological studies consistently show that bamboo dominance produces pronounced departures from woody evergreen broadleaf forest structure, including reduced tree basal area, altered gap dynamics, suppressed or redirected tree regeneration, and demographic processes driven by clonal expansion and episodic flowering–dieback cycles (Tabarelli and Mantovani 2000; Lima et al. 2012; Fadrique et al. 2021; Zheng and Lv 2023). These stands therefore represent fundamentally different canopy architectures, consistent with the structural contrasts observed in GEDI footprint-level LiDAR data (Fig. 3).

Mechanistically, bamboo differs from trees in ways that directly violate woody biomass allometries. Bamboo lacks secondary growth, forms hollow culms, and expands clonally through rhizomes (Makita 1998). Whereas tree biomass accumulates through continued radial growth of woody stems—supporting power-law relationships between DBH, height, and biomass (Enquist et al. 1999; Chave et al. 2014)—individual bamboo culms reach near-final height and diameter within a single growing season. Consequently, stand-level biomass increases primarily through culm recruitment and turnover rather than through growth of existing individuals (Yen 2016). Although culm-level DBH–height allometries exist (Sharma et al. 2025), estimating bamboo stand biomass requires explicit consideration of colony structure and culm age composition, complicating direct scaling from tree-based models (Zhang et al. 2014).

These biological differences intersect directly with GEDI’s biomass retrieval logic. GEDI Level 4A relies heavily on relative height (RH) metrics (e.g., RH50, RH98), implicitly assuming that waveform-derived vertical structure provides a consistent proxy for woody biomass across vegetation types (Kellner et al. 2023). In bamboo-dominated canopies, however, RH metrics may reflect dense foliage and hollow culm packing rather than woody stem mass, yielding an RH–AGBD relationship that deviates from the tree-calibrated EBT relationship assumed by GEDI L4A. Our height-stratified diagnostics (Fig. E.1; Method E) confirm that residual bias varies systematically with canopy height, indicating structural bias intrinsic to applying the EBT formulation to bamboo-dominated canopies.

Taken together, these considerations indicate that systematic bias is expected when GEDI’s global, RH-based EBT biomass model is applied to bamboo-dominated forests. Our analyses in Fig. 3 and Fig. 4 were designed to diagnose and quantify this bias, rather than to challenge the validity of GEDI products per se. Instead, our results align with the CEOS validation framework by emphasizing transparency in model assumptions and responsible interpretation of satellite-derived biomass estimates (Kay 2021). In bamboo-dominated forests, botanical traits such as clonal growth, hollow culms, and rapid culm maturation fundamentally shape canopy structure and biomass accumulation and must be considered when interpreting RH-based biomass products.

### 4.2 Scope, assumptions, and diagnostic interpretation

#### Diagnostic scope of the case study

This analysis is based on a single regional case study and is designed to develop a workflow for diagnosing and quantifying systematic bias in GEDI biomass estimates, rather than to provide definitive estimates of absolute biomass or carbon density differences between bamboo- and tree-dominated forests. Accordingly, the magnitude of bias we report should be interpreted as an order-of-magnitude reference rather than a universal correction factor. Our results are derived from one ecosystem and one broad contrast (bamboo versus evergreen broadleaf trees), and the magnitude of bias is therefore expected to vary across regions, species compositions, and successional contexts. Classification relies on single-year Sentinel-2 composites because of limited GEDI coverage in Xishuangbanna; incorporating time-series features would likely improve robustness by leveraging temporal patterns of phenological signals.

#### Benchmarking strategy and data constraints

A second limitation is the absence of site-specific ground-based biomass measurements in Xishuangbanna. While direct field measurements would provide the strongest benchmark, existing literature values are themselves uncertain. Our analysis therefore does not aim to establish a precise “true” biomass baseline. Instead, our conclusions rest on internally consistent comparisons among GEDI-derived structural metrics, GEDI’s operational EBT biomass model, and published empirical ranges (Yuen et al. 2017). The convergence of structural, ecological, and modeling evidence supports our central conclusion that applying tree-centric, RH-based biomass models to bamboo-dominated canopies introduces systematic bias.

#### Modeling assumptions and uncertainty treatment

Our uncertainty propagation relied on standard but simplifying assumptions. We treated the visually interpreted confusion matrix as an observation model and assumed that misclassification rates were stationary and that errors were conditionally independent given the mapped label. We used the same posterior resampling framework throughout the analysis: for footprint-level structural contrasts (Fig. 3; Table 1), we propagated uncertainty across all mapped classes to assess how tree–bamboo differences changed under plausible label reassignment; for the empirical-benchmark carbon comparison (Fig. 4; Table 2), we propagated uncertainty conditional on the mapped bamboo subset to maintain alignment with bamboo-specific benchmarks and context stratification (forest vs fallow). A fully landscape-level correction that also recovered omission errors (false negatives) would require reconstructing latent true-class composition across all land-cover classes and re-deriving bamboo context within each resampling iteration, which was beyond the scope of this workflow-focused case study.

Taken together, these limitations do not weaken our conclusions, because our objective is diagnostic rather than predictive: to identify the presence, direction, and structural origin of systematic bias in GEDI biomass estimates. Importantly, our contribution extends beyond bias identification. Using bamboo as a test case, we show that explicitly stratifying structurally distinct canopies using texture and structural features transforms the same remote-sensing data from a source of bias into an opportunity for refinement. This logic generalizes beyond bamboo to other vegetation types whose structure departs from global-model assumptions, including other grassy trees such as palms and bananas, and to fragmented agroforestry systems. We next outline applications that demonstrate how this workflow supports more responsible, bias-aware use of GEDI and related LiDAR products.

### 4.3 From regional case study to scalable global applications

Our workflow highlights the complementarity between horizontal information from optical imagery (spatial distribution and land-use context) and vertical information from LiDAR (canopy structure), both of which are necessary to interpret ecosystem function and avoid misrepresenting structurally distinct canopy types in biomass estimation. More broadly, stratification is a problem of scale selection under a coverage–accuracy trade-off: the features that best separate target vegetation classes often emerge at intermediate spatial or temporal scales, where signals can be detected consistently across heterogeneous landscapes. These results further emphasize that remote-sensing–derived products must be interpreted with biological awareness, because trait-driven deviations that are negligible at global scales can dominate local error budgets where those traits are prevalent.

One immediate application of this framework is place-based monitoring in structurally complex landscapes. By combining Sentinel-2 optical imagery with GEDI LiDAR structure, the workflow enables explicit identification and mapping of locally relevant canopy strata or species of interest, such as grassy trees (e.g., bamboo, palms, and bananas) that are fast-growing yet frequently conflated with trees in global products (Zheng and Lu 2025). This capability is particularly important in fragmented forests and agroforestry mosaics, where compositional changes occur over spatial scales too fine for coarse plant functional type classifications. Rather than relying solely on globally defined PFTs, the workflow allows locally meaningful strata to be detected using region-specific training data and local knowledge, supporting community-led monitoring in heterogeneous landscapes.

A second application lies in improving measurement, reporting, and verification (MRV) of forest carbon. Once structurally distinct vegetation types are explicitly identified, the same framework enables more targeted carbon accounting by reducing systematic bias that arises when biomass models are applied beyond their domain of transferability. This is especially relevant for REDD+ and other carbon-crediting or monitoring programs operating in secondary forests, forest–agriculture transition zones, and agroforestry systems, where grassy trees are often promoted or maintained through human management and can contribute substantially to landscape-level biomass yet remain poorly represented in global allometric frameworks (Tang et al. 2025). By separating bias driven by structural mismatch from true spatial variation in biomass, the workflow supports more transparent MRV.

Beyond carbon accounting, stratified structure–biomass estimates can guide targeted regeneration, restoration, and management interventions. In multilayered systems such as cocoa-and coffee-based agroforestry, LiDAR-enabled separation of canopy layers—combined with spectral or phenological indicators—can help identify vegetation components that contribute most to productivity, microclimate buffering, or climate resilience (Atalaya-Marin et al. 2025; Pippi et al. 2025). More broadly, our stratify–quantify–scale approach can be extended to track successional trajectories by separating shifts in canopy species composition from disturbance-driven changes in canopy structure, enabling interventions tailored to different successional stages.

## Conclusion

As remote sensing technologies advance and global products like GEDI become more accessible, they create new opportunities while also increasing the risk of misinterpretation. Responsible use therefore requires bias-aware interpretation grounded in ecology, organismal biology, land-use history, and local knowledge. Using bamboo as an example, we demonstrate that biologically informed canopy stratification can diagnose and quantify systematic bias in GEDI’s tree-calibrated biomass estimates and provide a practical workflow for doing so. Looking ahead, conceptual and technical advances can further improve the decision value of global products by specifying when stratification is needed, which signals reliably distinguish canopy types, and what validation is sufficient for targeted applications.

## Data Availability Statement

All remote-sensing datasets used in this study are publicly available. Data sources and processing details, together with all analysis scripts, are provided at https://github.com/aiyuz/GEDI_Carbon_Workflow.

## Author Contributions

AZ, YY, and ML jointly developed the conceptual framework of carbon discrepancy from structural heterogeneity. AZ implemented the workflow, analyzed the data, and prepared all figures and tables. AZ wrote the manuscript and compiled the scripts, with revisions from ML and YY.

## Acknowledgements

We thank Thaise Emilio, Ximena Londoño, Juan Carlos Camargo García, Junwei Luan, Cuiju Liu, Wenjun Zhou, and Jiayue Zhang for their collaboration in establishing the grassy tree research network. We also thank members of the Lu Lab and the Yin Lab in the Department of Environmental Studies at New York University for helpful discussions.

## Declaration of generative AI and AI-assisted technologies

During the preparation of this work, the authors used ChatGPT (version 5.2) to help organize and annotate scripts for publication and to check the language and grammar of the manuscript. After using this service, the authors reviewed and edited the content and take full responsibility for the content of the published article.

