## Appendix A-E for "Carbon bias from grassy tree misclassification: revealing forest structural heterogeneity by integrating GEDI and Sentinel-2"

Codes to reproduce results in this study are accessible at https://github.com/aiyuz/GEDI_Carbon_Workflow.

**Table of Contents**

**A** Bamboo-specific metrics and Gray-Level Co-occurrence Matrix of pixel intensities

- **Table A.1.** Visual interpretation clues used for manual labeling of land-use classes.
- **Table A.2.** Sample sizes of expert labeling and stratified value extraction.
- **Figure A.1.** Training labels of seven different classes and typical scenarios in Maxar submeter imagery for visual inspection.
- **Figure A.2.** Mean reflectance profiles of vegetation classes across Sentinel-2 spectral bands by season.
- **Figure A.3.** Spectral bands selected for classification and their distributions by class in training data.

**B** Classifier training and validation

**C** Propagating classification uncertainty in GEDI structural comparisons

- **Table C.1.** Confusion matrix from visual validation of the random-forest classifier (Google Earth, 2020–2022 imagery).
- **Figure C.1.** Coverage of GEDI LiDAR Level 4A aboveground biomass density footprints in Xishuangbanna (2021).
- **Figure C.2.** Misclassification-aware structural differences between bamboo-dominated and tree-dominated footprints.

**D** Empirical benchmarking for bamboo aboveground biomass density.

- **Table D.1.** Summary of aboveground carbon density data for bamboo from Yuen et al. (2017).

**E** Height-stratified evaluation of EBT-model residual bias

- **Figure E.1.** Height-stratified residual bias of the GEDI EBT biomass model for bamboo- and tree-dominated forests.

**Appendix A.** Bamboo-specific metrics and Gray-Level Co-occurrence Matrix of pixel intensities

**Table A.1** Visual interpretation clues used for manual labeling of land-use classes

| Class | Spatial Distribution | Texture | Seasonal Color Characteristics | Interpretation Basis |
|---|---|---|---|---|
| Rubber Plantations | Typically located on hillslopes along riverbanks, forming large continuous blocks | Densely aligned in rows, resembling contour lines in aerial view | Grey-yellow, light brown color in dry season or following leaf shredding or po | Recognizable plantation texture + known regional rubber distribution |
| Bamboo | Two main forms: (1) near villages, urban greenspaces, and field edges as planted clumping bamboo; (2) large patches along the Lancang River and in secondary forests | (1) Centrally spreading clumps, lacking individual tree crowns; (2) Strip-like canopy distinct from typical tree crowns | Lighter green in rainy season, turning yellow or pink in dry season | Species-specific morphology and ecological knowledge of bamboo distributions |
| Trees (non-bamboo forest) | Usually found in natural or secondary forests, without significant bamboo presence | Clear individual crowns, high heterogeneity in structure | Moderate seasonal color variation | Diverse crown shapes and canopy textures distinguish trees from bamboo |
| Plantation | Found on gentle slopes or hillsides, often small in area, used for fruit or other crops | Sometimes row-planted but with lower density, canopy not fully closed | Varies by crop type | Identified by lower crown closure and sparse structure |
| Farms | Located in flat areas or terraced valleys, mostly for grains and vegetables | Patchy, coarse texture, no woody canopy structure | Green during growing season; bare soil tones otherwise | Topographic position and lack of tree-like structures helps distinguish cropland |
| Construction | Located near roads or settlements | Sharp edges, high reflectance, clear building geometry | Gray or white tones; very bright in reflectance | Obvious anthropogenic structure in imagery |
| Water | Includes rivers, lakes, ponds, found in topographic lows | Smooth, texture-less, either very dark or highly reflective, depending on sun angle | Dark blue or black; spectral variability due to water depth | Reflectance and terrain context used to identify water bodies |

**Table A.2** Sample sizes of expert labeling and stratified value extraction.

| Class | Labeled Polygon (n) | Sampled Points (n) |
|---|---|---|
| Bamboo | 779 | 139767 |
| Construction | 37 | 31132 |
| Farms | 55 | 117479 |
| Plantation | 33 | 10825 |
| Rubber | 161 | 205922 |
| Trees | 176 | 236014 |
| Water | 29 | 41472 |

**Figure A.1** Training labels of seven different classes and typical scenarios in Maxar submeter imagery for visual inspection. a) The distribution of labeled training data with their elevation in Xishuangbanna. b) Landscape contexts for each label with details presented in Table A.1.

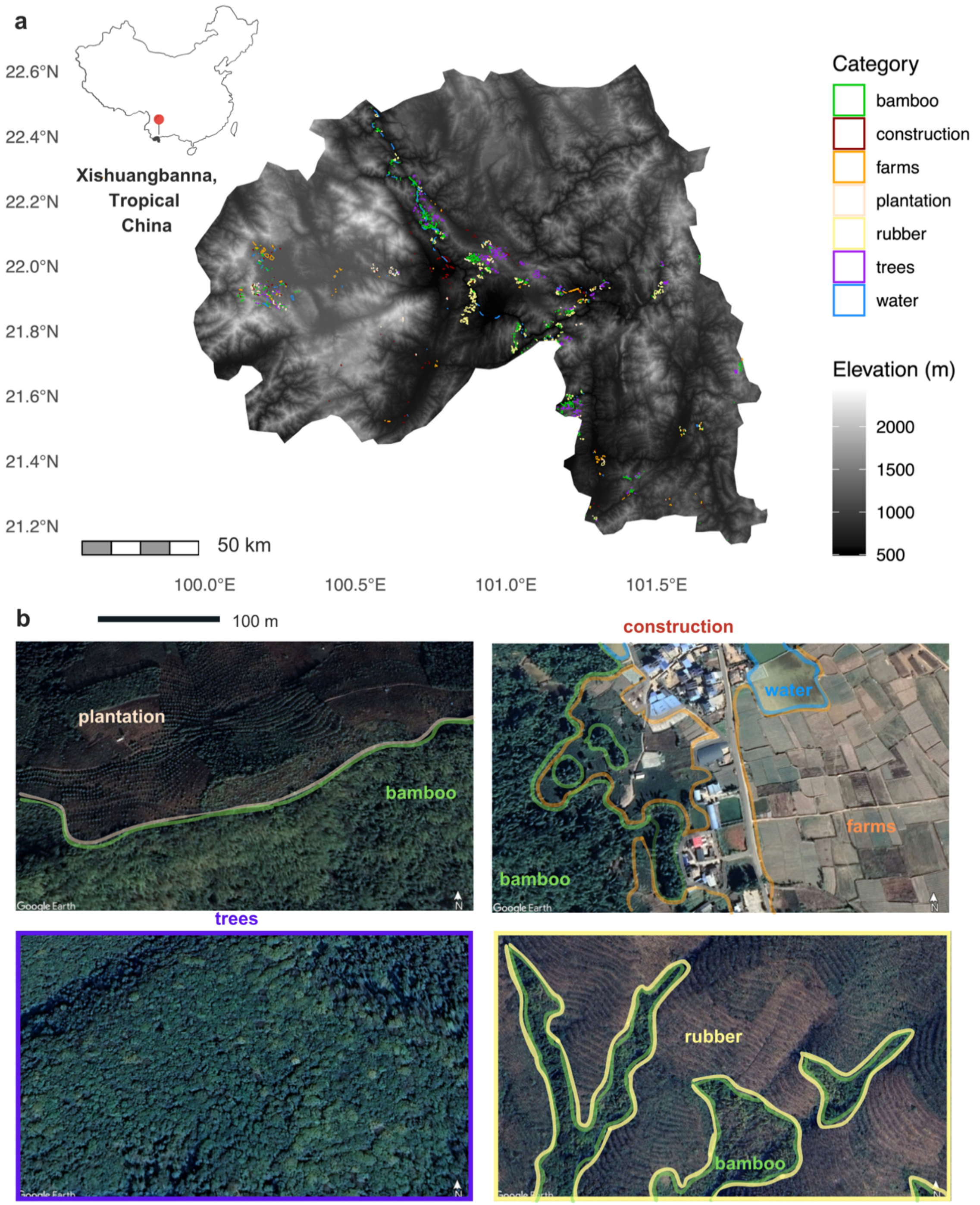

**Figure A.2** Mean reflectance profiles of vegetation classes across Sentinel-2 (Drusch et al. 2012) spectral bands by season. The July-September period didn't cover the entire study region; the October–September period was selected for compiling training data because it provides full coverage of Xishuangbanna, minimal cloud contamination, and relatively strong spectral separability. The NIR (Band 8) was chosen for texture analysis (following GLCM, see Haralick et al. 1973 and Mohammadpour et al. 2022) because it exhibits the greatest variance among the bands.

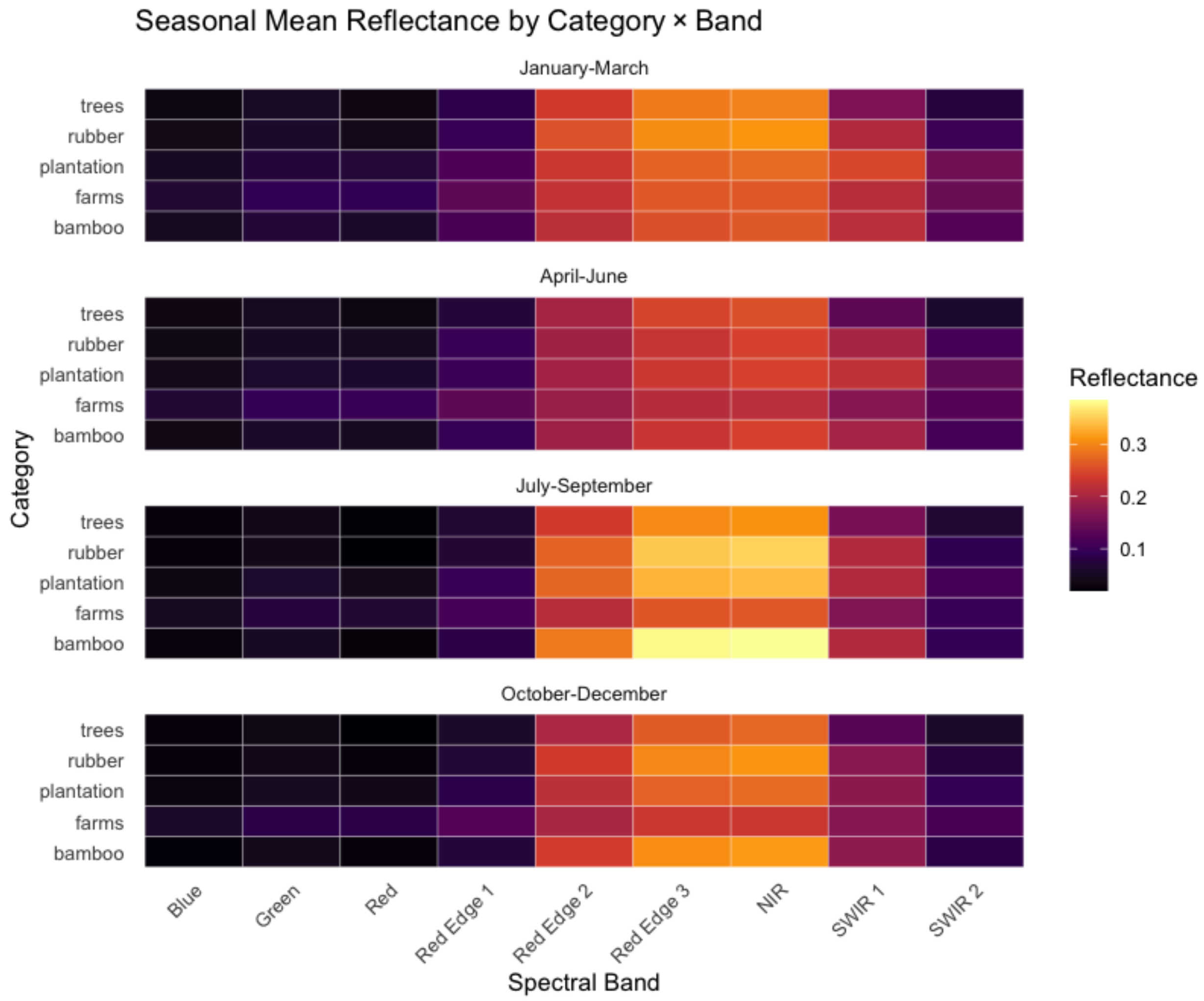

**Figure A.3** Spectral bands selected for classification and their distributions by class in training data.

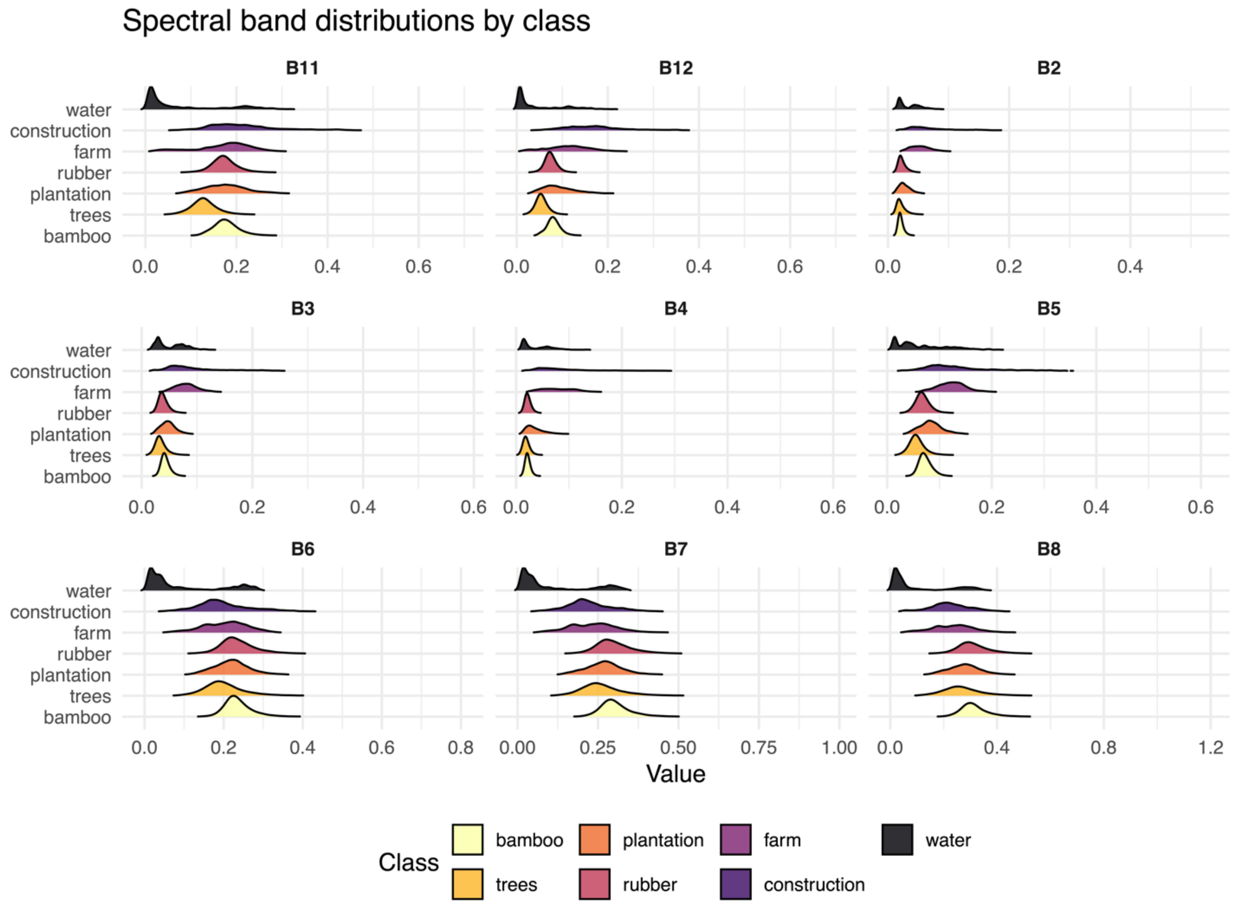

**Appendix B.** Classifier training and validation

In our study, we evaluated multiple supervised classifiers, including a random forest (RF), a one-dimensional convolutional neural network (1D-CNN), a fully connected artificial neural network (ANN), and a linear support vector machine (SVM), using the same pixel-level training dataset. Input features were derived from a 10-m resolution feature stack (October–December Sentinel-2 spectral bands, vegetation indices, texture metrics, and a canopy-height layer). Training samples extracted from labeled polygons were stratified by class and randomly split into 70% for model training and 30% for validation. Model performance was assessed using confusion matrices and overall accuracy.

The RF classifier was fitted using the randomForest package (v4.7-1.2) (ntree = 100; mtry = 4/5; importance = TRUE) (Liaw & Wiener 2002). Neural-network models were trained on z-score–standardized features. The 1D-CNN treated each pixel's feature vector as a one-dimensional sequence and was trained with a SoftMax output layer, Adam optimizer, and categorical cross-entropy loss (maximum 50 epochs, batch size = 32, early stopping with patience = 5). The ANN employed two dense hidden layers with dropout and the same optimization and stopping criteria. The SVM used a linear kernel with feature centering and scaling implemented via caret.

Based on this initial comparison of model performance on the held-out validation set, we selected the two best-performing models (RF and 1D-CNN) for further independent visual validation using high-resolution Google Earth imagery. This additional step was used to assess class-specific sensitivity and specificity beyond the training/validation data. The RF classifier showed the most consistent performance and spatial stability in this external validation and was therefore used for all final 10-m land-cover maps and subsequent analyses involving GEDI data in the main study.

**Appendix C.** Propagating classification uncertainty in GEDI structural comparisons

**Bayesian updates on land class probabilities based on their misclassification rates**

*Misclassification rates*

Each land use class in the classified map was randomly sampled for 100 spatial points for estimating misclassification errors. The coordinates of sampled points were imported into Google Earth Pro (version 7.3.6, Google Earth 2025) to examine whether its classified class was correctly identified using high-resolution maps during 2020-2022, depending on the availability of imagery for each tile within the study region.

If the actual class of a point was bamboo while it was misidentified as any other class, the case was considered a false negative. If the actual class of a point was not bamboo while it was identified as bamboo, the case was considered a false positive. The false positive and false negative rates of each class in 2021 were summarized in Table C.1. The misclassification information can be stored in a confusion matrix for calculating the probability of a pixel's true class in a way similar to Bayesian updating.

We define a few terms here:

1) The true probability of a pixel being in class $k$ is $Q_k$.
2) Every pixel in the study region has probability $p_k$ to be identified by the classifier as class $k$.
3) As we obtain information from the classifier initially, $p_k$ is the ratio of pixels identified as $k$ to total number of pixels, where $m$ is the total number of classes.

$$p_k = \frac{N_k}{N_{total}} \text{ and } \sum_{k=1}^{m} p_k = 1 \text{ (Eq. C1)}$$

4) Now the validation information in the confusion matrix can update our initial plausibility, $p_k$.

For a five-class classifier, $\left(C_{i,j}\right)_{i,j\in\{1,\dots,5\}}$ is its confusion matrix where the rows indicate results from the classifiers and the columns indicate true categories. $C_{i,j}$ represents the number of cases for actual class $j$ being identified as class $i$.

5) The misclassification likelihood of a pixel identified as class $i$ and validated to be in class $j$ is denoted as $L_{i,j}$. When $i = j$, the classifier accurately predicted the class. The sample size of validation data for each class $i$ is the sum of entries in row $i$ of the confusion matrix. Therefore:

$$L_{i,j} = \frac{C_{i,j}}{\sum_{j=1}^{m} C_{i,j}} (Eq. C2)$$

The confusion matrix tells us that the probability of a pixel being a true class $j$ is the probability of such pixel identified as class $j$ given a correct identification, plus the sum of probabilities of the pixel being misidentified for class $i$ times the probability of a false negative report on class $i$ pixels. The joint probability of class $k$ is $P_k$:

$$P_k(classifer, validated) = \sum_{j=1}^{m} p_i \times L_{i,j=k} (Eq. C3)$$

$$P_k(validated) = \sum_{k=1}^{m} \sum_{i=1}^{m} p_i \times L_{i,j=k} (Eq. C4)$$

And thus, we can update the initial probability from classification, and let's denote this updated probability as $Q_k$:

$$Q_k(classifer \mid validated) = \frac{prior \times probability\ of\ observations}{Average\ probability\ of\ observations}$$
$$= \frac{\sum_{j=1}^{m} p_i \times L_{i,j=k}}{\sum_{k=1}^{m} \sum_{i=1}^{m} p_i \times L_{i,j=k}} (Eq. C5)$$

Using steps 1-5 described above, we updated the probability of each class type to calculate the total bamboo-dominated area in Xishuangbanna. Confusion matrices acquired from visual accuracy assessment (photo-interpretation-based validation) were included in Table A.1. The overall accuracy was determined by the fraction of correctly identified cases (sum of the diagonal elements in a confusion matrix) to total cases identified (sum of all elements in a confusion matrix). The user's accuracy for class $k$ is the fraction of correctly identified $k$-class cases to total identified cases for class $k$. The producer's accuracy is the fraction of correctly identified cases in class $k$ to all cases that are validated as class $k$.

*Confusion Matrix*

The confusion matrix was acquired by visually examining classification results via the random forest model. The overall accuracy was 84.9%. For bamboo, containing bamboo-dominated forest areas and bamboo patches scattered throughout the landscape, the user accuracy and producer accuracy were 92% and 90.2%, respectively. For trees, which contain extensive preserved forest area and forest remnants under more frequent human disturbance, the user accuracy and producer accuracy were 92% and 82.1%, respectively.

**Table C.1** Confusion matrix from visual validation of the random-forest classifier (Google Earth, 2020–2022 imagery).

| Classified class ($i$) | Validated class ($j$) | | | | | | |
|---|---|---|---|---|---|---|---|
| | B | C | F | P | R | T | W |
| Bamboo | 92 | 1 | 1 | 2 | 4 | 6 | 0 |
| Construction | 0 | 76 | 3 | 0 | 0 | 0 | 5 |
| Farms (Crop) | 0 | 14 | 88 | 7 | 0 | 0 | 2 |
| Plantation (Fruit/Tea) | 3 | 7 | 8 | 78 | 5 | 0 | 0 |
| Rubber | 5 | 2 | 0 | 9 | 76 | 2 | 0 |
| Trees (Forest) | 0 | 0 | 0 | 4 | 15 | 92 | 1 |
| Water | 0 | 0 | 0 | 0 | 0 | 0 | 92 |

**Figure C.1** Coverage of GEDI LiDAR Level 4A aboveground biomass density footprints in Xishuangbanna (2021), overlaid on 10-m bamboo and tree classifications derived from the random-forest classifier.

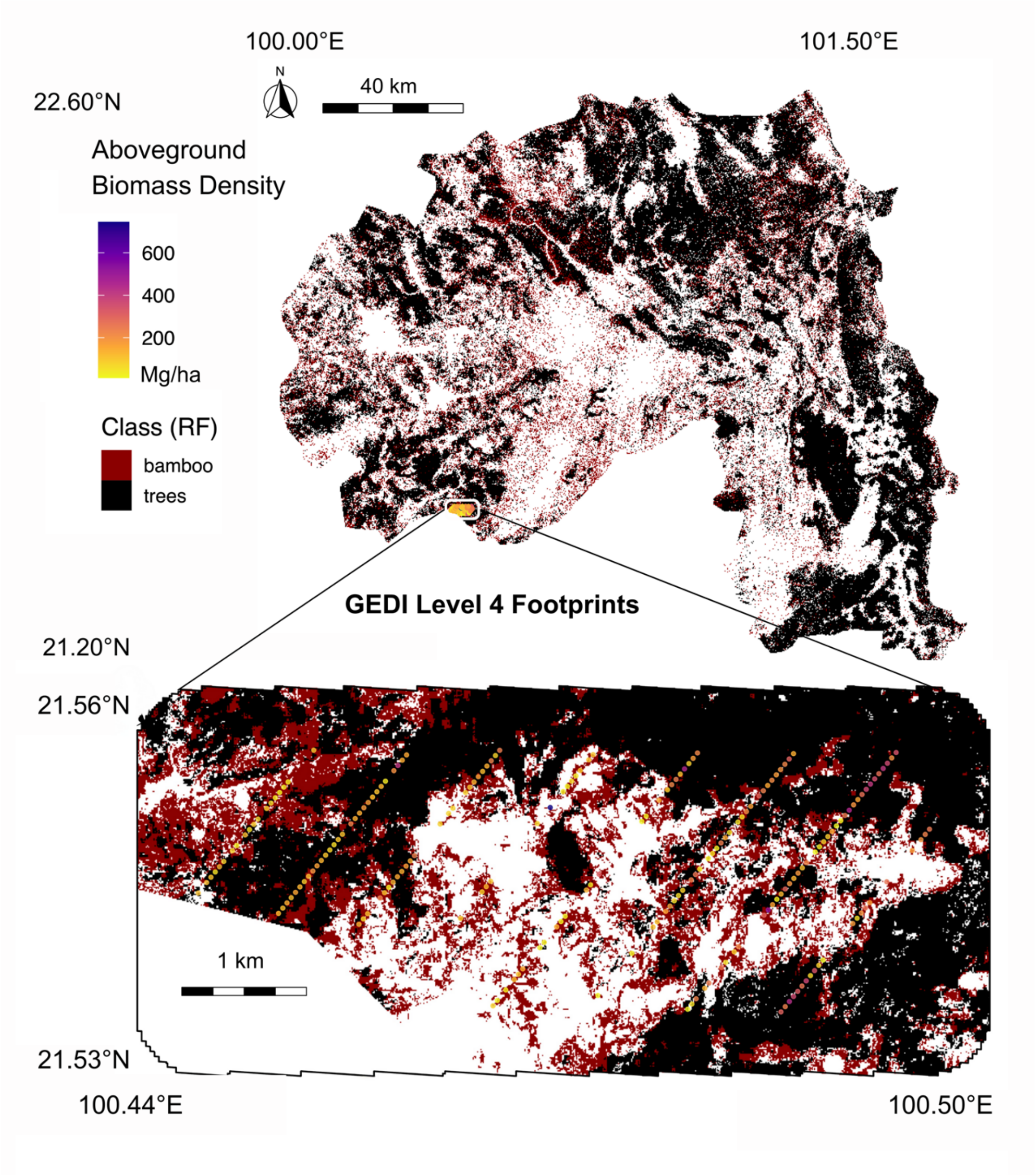

To account for land-cover misclassification when comparing GEDI-derived structural metrics between bamboo and tree-dominated forests, we propagated classification uncertainty using a Bayesian posterior resampling approach based on the externally validated confusion matrix described in Appendix C and Table C.1.

Briefly, we derived the likelihood matrix $P(predicted \mid actual)$ from the visually validated confusion matrix and estimated class priors $P(actual)$ from pixel-wise class frequencies in the regional Random Forest classification map. These components were combined to obtain posterior class probabilities $P(actual \mid predicted)$ for each predicted land-cover class.

For each GEDI footprint and structural metric (canopy height, aboveground biomass density, plant area index, and total vegetation volume), we performed Monte Carlo resampling (n = 1000 iterations), in which footprint-level observations were reassigned to a "true" land-cover class by drawing from the corresponding posterior distribution. Median differences between trees and bamboo were re-estimated in each iteration using Wilcoxon rank-sum tests.

Across all structural metrics, posterior-corrected median differences remained consistently positive, and the proportion of iterations with $p < 0.05$ was high (often equal to 1.0), indicating that the observed bamboo–tree contrasts are robust to plausible land-cover misclassification errors.

**Figure C.2** Misclassification-aware structural differences between bamboo-dominated and tree-dominated footprints.

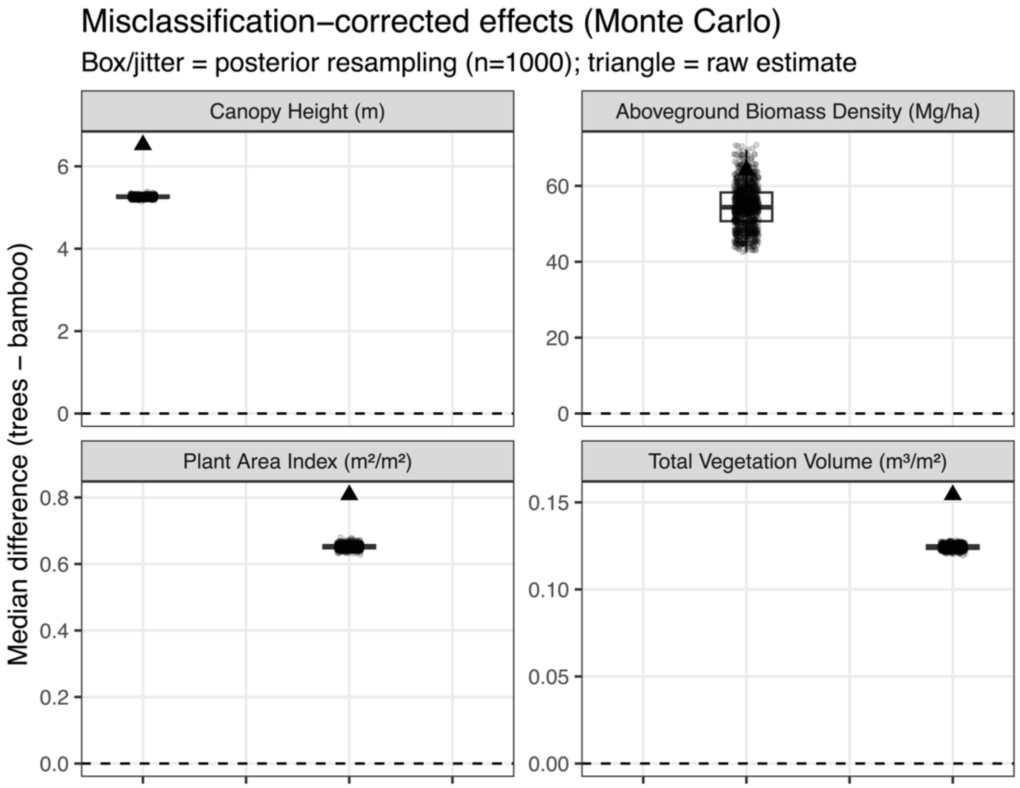


**Appendix D.** Empirical benchmarking for bamboo aboveground biomass density.

**Derivation of confidence intervals from literature summary statistics**

Empirical benchmarks for bamboo aboveground carbon density were obtained from Yuen et al. (2017), which reports summary statistics including sample size ($n$), sample mean ($\underline{x}$), sample standard deviation ($s$), and maximum observed values for multiple bamboo types and contexts (see Table D.1). Because individual-level observations were not available, confidence intervals for the empirical mean carbon density were reconstructed from these published summary statistics following standard statistical practice.

**Assumptions**

Our reconstruction assumes that observations within each bamboo context represent independent samples from an underlying population, and that the sampling distribution of the mean is approximately normal. This assumption is justified by the Central Limit Theorem for moderate sample sizes and is standard for inference on population means when only summary statistics are available.

**Confidence interval for the mean**

For each bamboo context used in our analysis, the 95% confidence interval (CI) of the population mean aboveground carbon density was computed using the *t*-distribution:

$$\underline{x} \pm t_{0.975,n-1} \times \frac{s}{\sqrt{n}} (Eq. D1)$$

Where $\underline{x}$ is the reported sample mean, $s$ is the reported sample standard deviation, $n$ is the sample size, and $t_{0.975,n-1}$ is the 97.5th percentile of the *t*-distribution with n−1 degrees of freedom. This formulation accounts for uncertainty in the estimated variance and is appropriate when population variance is unknown.

**Use of maximum values**

Maximum values reported in Yuen et al. (2017) were not used in the calculation of confidence intervals. Instead, they were retained as reference points to contextualize the empirical range of observed bamboo carbon densities. In Fig.4a, these maxima are shown as dashed horizontal lines to illustrate how GEDI-derived estimates compare not only to the empirical mean and its associated uncertainty, but also to the upper bounds reported in field measurements.

**Interpretation**

The reconstructed confidence intervals quantify uncertainty around the empirical mean bamboo carbon density, rather than the full spread of observed values. Accordingly, GEDI-derived estimates exceeding these confidence intervals indicate systematic overestimation relative to the mean field-based benchmark, even after accounting for sampling variability in the original empirical study.

**Table D.1** Summary of aboveground carbon density data for bamboo from (Yuen et al. 2017). The first two rows were used as regional empirical benchmarks in our analysis; other entries are included for context because they represent different species, sites, or measurement contexts and were not used in statistical tests.

| Species Name | Location | Sample size | Mean AGC | SD AGC | Max AGC |
|---|---|---|---|---|---|
| Bamboo fallow | India, Laos, Myanmar | 35 | 14.7 | 14.1 | 56.4 |
| Bamboo forest | China, Laos, Myanmar, Thailand, Vietnam | 24 | 27.5 | 43.1 | 162.0 |
| *Bambusa arudinacea* | India | 6 | 23.5 | 17.9 | 50.9 |
| *Bambusa bambos* | India | 13 | 81.1 | 46.0 | 143.3 |
| *Bambusa oldhami* | China | 9 | 25.7 | 27.7 | 71.6 |
| *Bambusa polymorphia* | Myanmar | 13 | 15.3 | 9.1 | 31.8 |
| *Bambusa tulda* | Bangladesh, India, Myanmar, Philippines | 11 | 23.5 | 17.0 | 53.0 |
| *Dendrocalamus giganteus* | China | 6 | 33.6 | 36.4 | 77.9 |
| *Dendrocalamus latiflorus* | China | 22 | 15.3 | 15.7 | 57.0 |
| *Dendrocalamus strictus* | China | 8 | 20.7 | 15.5 | 49.1 |
| *Gigantochloa spp* | Indonesia, Thailand | 4 | 23 | 15.4 | 43.7 |
| *Thyrsostachys siamensis* | Thailand | 4 | 17 | 9 | 26.9 |

**Appendix E.** Height-stratified evaluation of EBT-model residual bias

To examine how bias in the simplified GEDI evergreen broadleaf tree (EBT) biomass model varies with canopy height, we analyzed residuals of aboveground biomass density (AGBD) as a function of RH98, separately for bamboo- and tree-dominated footprints.

For each GEDI footprint, residual AGBD was defined as the difference between biomass predicted from the simplified EBT formulation using canopy-derived RH98 and RH50 ($AGBD_{EBT}$) and the corresponding GEDI Level 4A estimate ($AGBD_{L4A}$):

$$Residual\ AGBD_i = AGBD_{EBT,i} - AGBD_{L4A,i} (Eq. E1)$$

Footprints were grouped into discrete RH98 height bins (10–20 m, 20–30 m, 30–40 m, and 40–50 m, where available) to evaluate how residual bias varies across canopy height classes. Bamboo-dominated footprints were limited to bins where sufficient observations were available (10–30 m), whereas tree-dominated footprints spanned a wider height range.

Within each RH98 bin and vegetation class, residual distributions were summarized using boxplots showing the median, interquartile range, and range excluding extreme outliers. A horizontal reference line at zero indicates unbiased agreement between EBT-predicted and GEDI L4A biomass.

This height-stratified analysis allows us to assess whether systematic over- or underestimation by the EBT model depends on canopy stature, and whether such patterns differ between bamboo- and tree-dominated systems. Because this comparison relies solely on GEDI-derived structural metrics and the operational EBT formulation, it isolates model-structural effects independent of external empirical benchmarks.

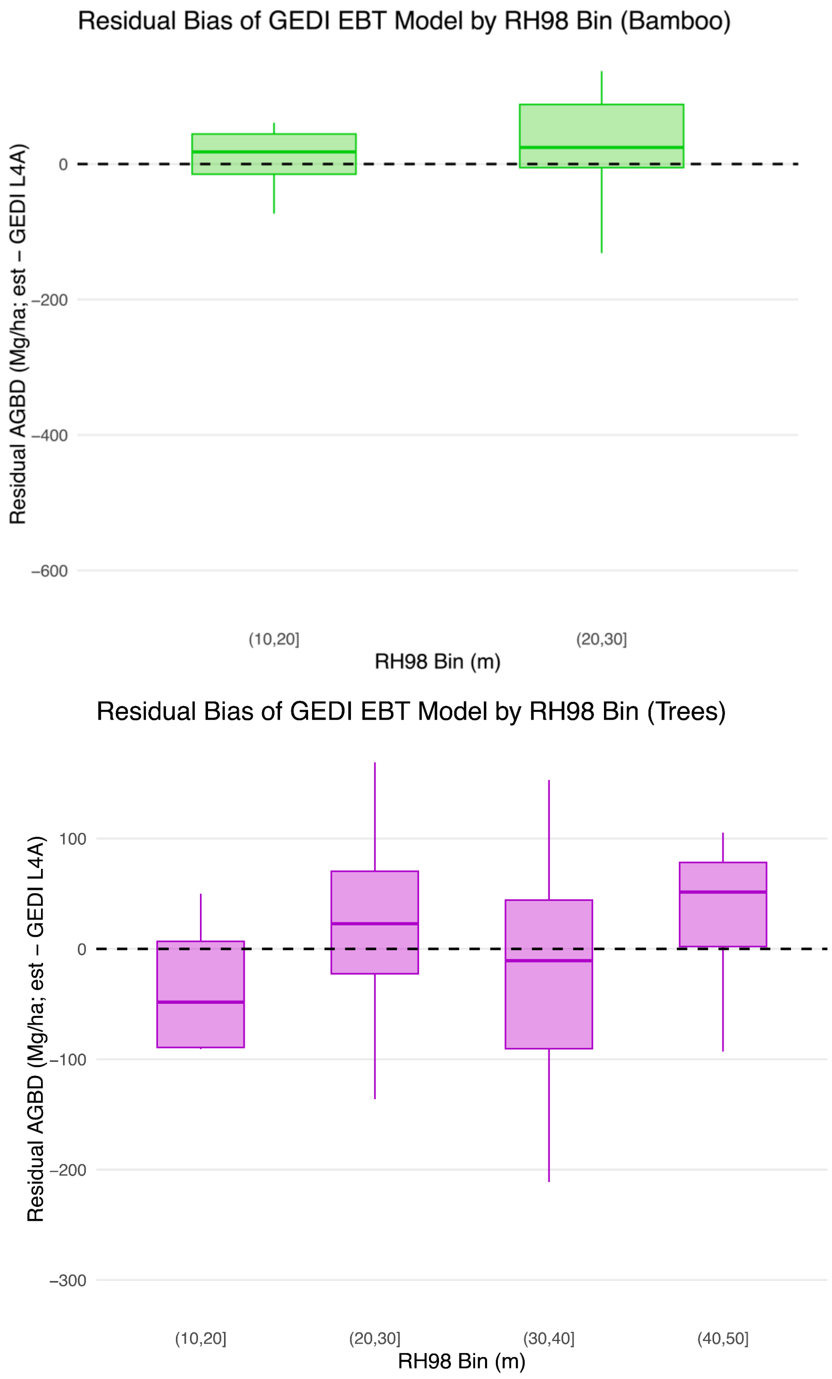


**Figure E.1.** Height-stratified residual bias of the GEDI EBT biomass model for bamboo- and tree-dominated forests. Positive residuals indicate overestimation by the EBT model relative to GEDI L4A, while negative values indicate underestimation. The contrasting patterns between bamboo and trees across height bins illustrate height-dependent and vegetation-type-specific bias arising from applying a single tree-based biomass model to structurally distinct forest systems.
